# Single cell view of tumor microenvironment gradients in pleural mesothelioma

**DOI:** 10.1101/2024.03.14.585048

**Authors:** Bruno Giotti, Komal Dolasia, William Zhao, Peiwen Cai, Robert Sweeney, Elliot Merritt, Evgeny Kiner, Grace Kim, Atharva Bhagwat, Samarth Hegde, Bailey Fitzgerald, Sanjana Shroff, Travis Dawson, Monica Garcia-barros, Jamshid Abdul-ghafar, Rachel Chen, Sacha Gnjatic, Alan Soto, Rachel Brody, Seunghee Kim-Schulze, Zhihong Chen, Kristin G. Beaumont, Miriam Merad, Raja Flores, Robert Sebra, Amir Horowitz, Thomas U Marron, Anna Tocheva, Andrea Wolf, Alexander M. Tsankov

## Abstract

Immunotherapies have shown great promise in pleural mesothelioma (PM), yet most patients still do not achieve significant clinical response, highlighting the importance of improving understanding of the tumor microenvironment (TME). Here, we utilized high-throughput, single-cell RNA-sequencing to *de novo* identify 54 expression programs and construct a comprehensive cellular catalogue of the PM TME. We found four cancer-intrinsic programs associated with poor disease outcome and a novel fetal-like, endothelial cell population that likely responds to VEGF signaling and promotes angiogenesis. Throughout cellular compartments, we observe substantial difference in the TME associated with a cancer-intrinsic sarcomatoid signature, including enrichment in fetal-like endothelial cells, CXCL9+ macrophages, cytotoxic, exhausted, and regulatory T cells, which we validated using imaging and bulk deconvolution analyses on two independent cohorts. Finally, we show, both computationally and experimentally, that NKG2A-HLA-E interaction between NK and tumor cells represents an important new therapeutic axis in PM, especially for epithelioid cases.

**Statement of Significance:** This manuscript presents the first single-cell RNA-sequencing atlas of pleural mesothelioma (PM) tumor microenvironment. Findings of translational relevance, validated experimentally and using independent bulk cohorts, include identification of gene programs predictive of survival, a fetal-like endothelial cell population, and NKG2A blockade as a promising new immunotherapeutic intervention in PM.

## Introduction

Pleural mesothelioma (PM) is a cancer of the lung pleura that is strongly associated with exposure to asbestos (1), although the proportion of patients without known occupational asbestos exposure is rising (2). Histologic subtypes can be characterized as epithelioid (60-75% of cases), sarcomatoid (10%), or biphasic PM (20-30%), with the latter thought to represent a mixture of epithelioid and sarcomatoid subtypes (3). Due to the aggressive nature of all histological types (4,5), existing therapeutic strategies (6,7) have had limited success with a median overall survival of approximately 18 months (6). Recently, the combination of anti-PD1 and anti-CTLA4 checkpoint inhibitors has emerged as an effective combination therapeutic option for PM; despite similar response rates compared to chemotherapy, responses are more durable in this immunotherapy combination, resulting in a 27% decrease in the risk of death (8). Patients with sarcomatoid and biphasic (non-epithelioid) histologies have been historically associated with worst overall survival but are also marked by a higher lymphocyte infiltration in the tumor microenvironment (TME) (9) and show greater benefit from checkpoint blockade combination treatments relative to chemotherapy, which, in contrast, has greater efficacy in epithelioid tumors (8).

While immunotherapy holds great promise, most patients with PM still do not achieve significant clinical benefit from these therapies, and many who do respond initially only receive a transient benefit. Given the variability in response encountered among patients and the toxicities associated with these therapies, new approaches are needed to determine which patients will benefit from existing immunotherapies and to discover new therapeutic strategies for non-responders. It is likely that intra- and inter-tumoral heterogeneity in the TME and tumor-immune cell interactions all contribute to the variability in treatment response. Thus, a more complete characterization of the PM TME at baseline will reveal more optimal patient stratification strategies and new immunomodulatory pathways to target.

Large scale bulk genomic and transcriptomic studies (10–13) have defined molecular subtypes associated with differences in the TME composition, including higher levels of T cells and M2-like macrophages in sarcomatoid and enhanced VISTA expression in epithelioid PM (10,11). A following meta-analysis study further reported on higher lymphocyte and monocyte infiltration, increased stromal components and expression of immune checkpoints molecules in PM samples correlated with a sarcomatoid transcriptional phenotype (S score) whereas VISTA and natural killer (NK) cell markers trended in the opposite direction (14). Additional studies expanded on the PM tumor-subtype dichotomy to define novel subtypes based on additional molecular features such as immune content, DNA methylation and tumor ploidy (12,13). Similarly, a recent mass cytometry study based on a 35 antibody panel also identified two histology-independent immunologic subtypes related to MHC-I and MHC-II neopeptide abundance (15). Single-cell RNA-sequencing (scRNA-seq) now enables interrogation of the TME at unprecedented resolution and scale without *a priori* knowledge and reliance on a limited set of markers, which has greatly enhanced our understanding of tumor heterogeneity across cancers (16). Here, we used high-throughput, scRNA-seq and single-cell T Cell Receptor sequencing (scTCR-seq) on treatment naïve patient samples to build a comprehensive single-cell atlas of PM primary tumor and peripheral blood. Our integrative analysis allowed us to ask (1) if there are cellular and molecular differences in the TME between PM histological and molecular subtypes, (2) if different subtypes associate with different cancer-intrinsic programs, (3) if new cell type specific signatures are predictive of disease outcome, and (4) if the scRNA-seq data suggest more effective, personalized therapies.

## RESULTS

### A single-cell catalogue of patient-matched PM tumors and peripheral blood

Our study group included 13 treatment-naïve patients diagnosed with PM spanning all three histological subtypes and comprising of 4 non-white and 5 females patients (31% and 38% of total cohort respectively), providing greater diversity compared to national incidence demographics (17) (Table S1). Primary tumor samples were obtained either during surgical resections (n=7) or diagnostic biopsies (n=6) and profiled for scRNA-seq using the 10X Chromium platform, including scTCR-seq on 7 samples (Figure 1A). In parallel, peripheral blood mononuclear cells (PBMC) were similarly profiled for a subset of patients (n=8). Following stringent quality control, a total of 141,219 cells were recovered (Figure S1A; Methods). We constructed an analytical pipeline aimed at uncovering axes of molecular variation across cellular compartments and PM subtypes in our single cell data (discovery cohort), which we validated experimentally and *in silico* using bulk RNA-seq and patient survival data (Figure 1B) from Bueno et al., and Hmeljak et al., (10,11) (293 patients in total; hereafter named Bueno and Hmeljak validation cohorts). Unsupervised dimensionality reduction and clustering of the tumor scRNA-seq data (Figure 1C, left panels) allowed for unbiased discovery of both established and previously unreported PM markers (Figure 1D, Table S2) for all major cell types detected in the tumor samples, including tumor cells (*KRT19*), normal mesothelial cells (*HP*), fibroblasts (*COL1A1*), smooth muscle cells (*MYH11*), endothelial cells (*PECAM1*), myeloid (*LYZ*), T cells (*CD3D*), NK (*GNLY*) B cells (*CD79A*), plasma cells (*IGLC2*), plasmacytoid dendritic cells (pDC, *IRF8*), a small number of alveolar type II cells (AT2; *SFTPC*) and a rare glial population (*PMP2*) recovered in only one of the patients. Similarly, we identified transcription factors (TFs) most specifically expressed in each major cell type (Figure S1B), which agreed with the TFs’ known role, including TEAD1 in malignant cells, WT1 in mesothelium, and SNAI2 in fibroblasts (18–20). As samples were collected with two different procedures (biopsy or surgical resection) we examined difference in cell proportions which showed higher fractions of B, T, and NK cells in the resection samples and malignant cells in the biopsies (Figure S1C). To normalize for sample acquisition differences in cell composition, we performed downstream cell subset and program enrichment analyses relative to each cellular compartment and validated our main findings throughout the study with bulk deconvolution analysis.

**Figure 1.**
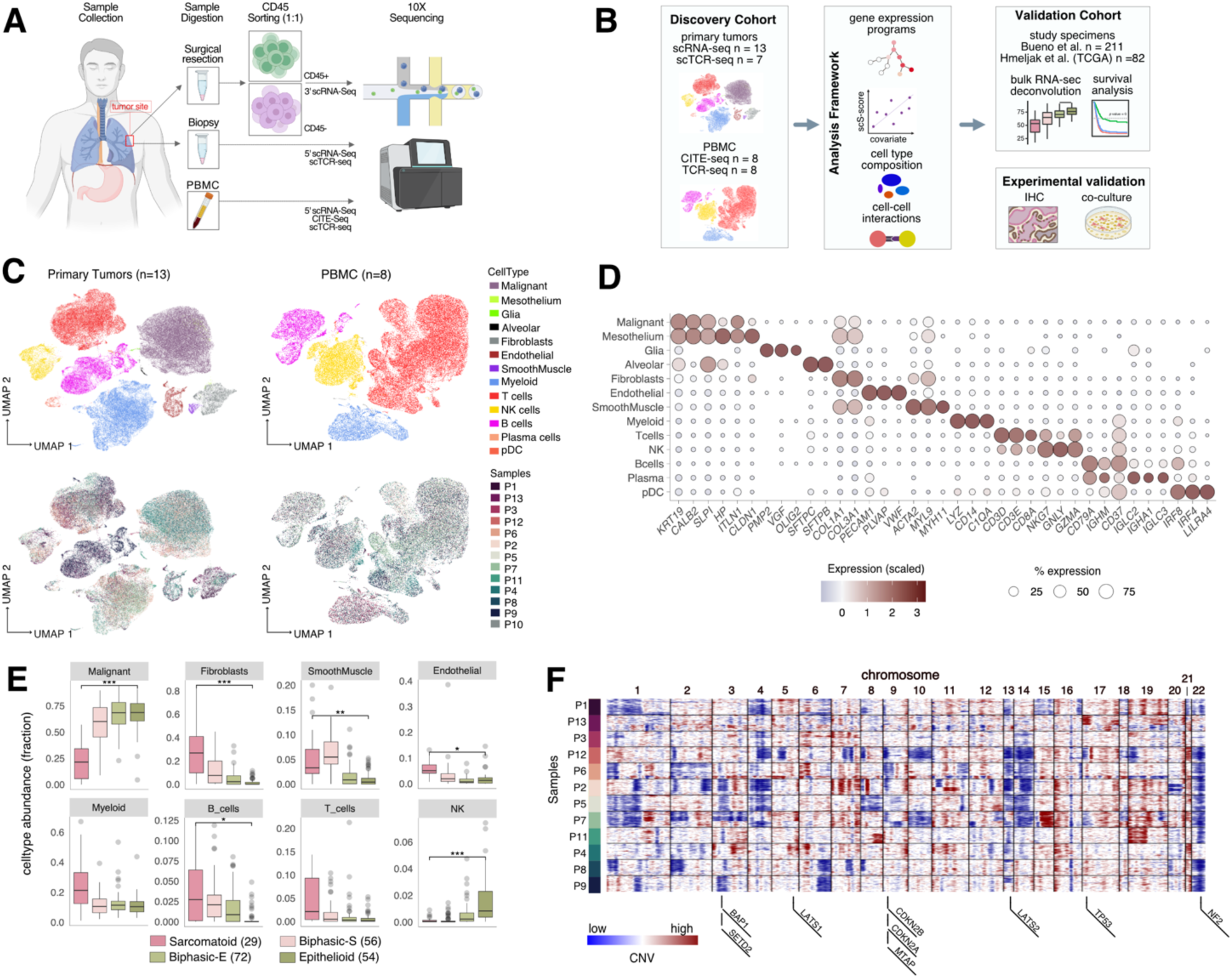
Single-cell catalogue of PM tumor and PBMC samples shows distinct TME cell composition differences between molecular subtypes. **A**, Schematic of sample collection, digestion, cell sorting, and sequencing. **B** Analysis workflow **C**, Uniform Manifold Approximation and Projection (UMAP) plots colored by cell type annotations (top) and patient identities (bottom) of primary tumors (left) and PBMC (right). **D**, Dot plot showing expression and percentage of cells expressing selected marker genes for each annotated cell type. **E**, Sample distributions based on cell type proportions (deconvolved using BayesPrism) in the Bueno cohort, grouped by different molecular subtypes. FDR-adjusted *P* values comparing difference between sarcomatoid and epithelioid subtypes were determined by Dirichlet-multinomial regression model that takes into account dependencies in proportions between cell types. *p<0.05, **p<0.01, ***p<0.001, ****p<0.0001. F, Inferred CNVs of malignant cells from primary tumor samples (sub-sampled to 200 cells per patient) with genomic location of key PM driver mutation genes shown on the bottom.

Additionally, we performed cellular indexing of transcriptomes and epitopes (21) (CITE-seq) to construct a patient-matched single cell atlas of the cellular protein and 5’ transcriptomes of PM PBMCs (Figure 1C, right panels). 30 PBMC subsets shared across the 8 patients were annotated using a reference-based pipeline (22), and *de novo* protein and RNA marker discovery identified canonical genes associated with these PBMC annotations (22), highlighting the quality of the data generated (Figure S1D-G; Tables S3-4).

We next investigated cell type abundance differences across PM molecular and histological subtypes using a Bayesian deconvolution framework powered by our PM-specific single-cell expression data (23). To robustly assess these data, we leveraged both the Bueno and Hmeljak validation cohorts. Results broadly agreed with previous bulk deconvolution cell type estimations (13,14), showing a more prominent infiltrate of T and B lymphocytes and myeloid immune populations as well as a more abundant stromal component in non-epithelioid subtypes, whereas epithelioid tumors were comparatively enriched in malignant and NK cells (12–14) (Figure 1E, S1H-I).

Inference of copy number variations (CNVs) enabled us to distinguish malignant cells from normal mesothelial cells in the tumor scRNA-seq data (Figure 1F), where we detected no malignant cells in biopsy sample P10 and therefore excluded it from all cancer cell downstream analyses. The CNV analysis detected large scale deletions on chromosomes 3 (p-arm), 13, 14 and 22 in most samples, in agreement with frequently deleted regions in PM detected by DNA sequencing (10,24), which harbor commonly deleted genes such as BAP1, LATS2, and NF2 (Figure 1F). Taken together, we have constructed the first comprehensive single cell atlas of PM and observe clear differences in TME cell type compositions between PM molecular and histological subtypes.

### *De novo* discovery of PM cancer programs show link to disease outcome

We reasoned that analysis of scRNA-seq data from 30,318 PM malignant cells can provide new, higher-resolution insight on intra- and inter-tumor heterogeneity. Towards this goal, we scored each malignant cell using signatures derived from four previously identified PM molecular subtypes (11) — sarcomatoid, biphasic-S, biphasic-E, and epithelioid (Figure 2A, left panel). As expected, we observed that in the most sarcomatoid (e.g., P1, P13) and epithelioid (e.g., P8, P9) tumors, malignant cells predominantly reside in the corresponding subtype quadrants (Figures S2A, 2A, right panel). However, several patients’ tumors histologically classified as predominantly epithelioid (e.g., P2, P7) were comprised of malignant cells that spanned all 4 molecular subtypes, uncovering a previously unappreciated intra-tumoral heterogeneity (Figures S2A, 2A, Table S1). Taken together, our data supports the view that PM tumors lie on a continuous spectrum between sarcomatoid and epithelioid subtypes (13,14) and further provides evidence that this paradigm is also valid at single-cell resolution.

**Figure 2.**
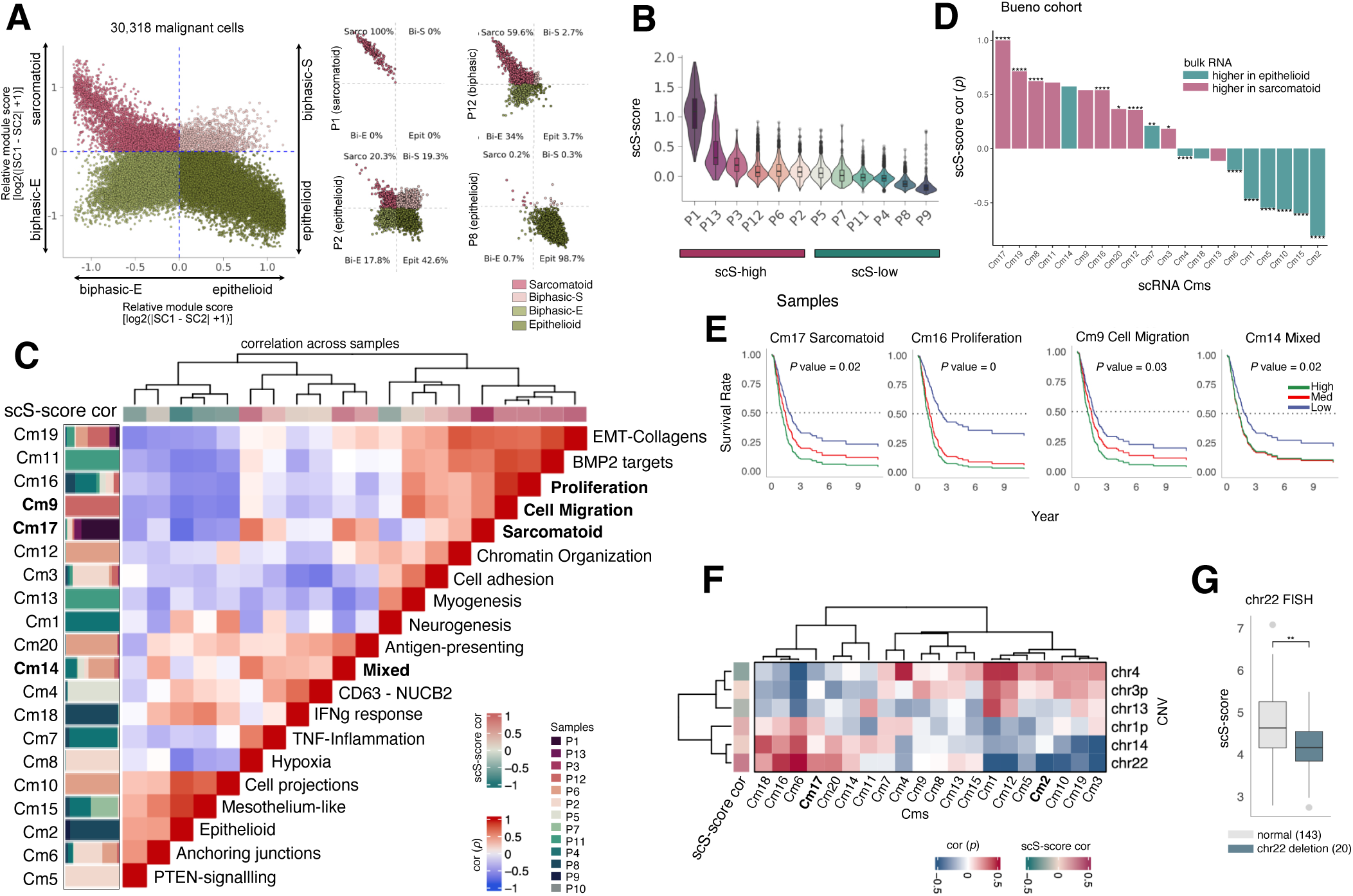
Unbiased discovery of PM cancer programs and association with patient survival. **A**, Left: Two dimensional (2D) representation (Methods) of the malignant cell distribution across the four PM molecular subtypes (quadrants) defined in Bueno et al (11) combining cells from all PM patients. Right: 2D representation of the malignant cell distribution for only four representative PM patients. Clinical histology of the cancers upon diagnosis are reported in parentheses. **B**, Per sample malignant cell distribution of the scS-score based on genes in Cm17, identified *de novo* from the scRNA-seq data. Tumor samples were classified as scS-high or scS-low based on their mean scS-score ranking. **C,** Pairwise Spearman correlation of sample-averaged scores derived from the 20 Cms identified in malignant cells. Each Cm was annotated with the most representative biological pathway. Vertical stacked bar plots (left) show Cm sample distribution. Top color bar shows correlation of each Cm to the scS-score **D**, Cm ranked by correlation to the sSC-score and colored by their enrichment for either epithelioid (green) or sarcomatoid (red) molecular subtypes from the Bueno cohort. FDR-adjusted *P* values were computed using Welch Two Sample *t*-test. *p<0.05, **p<0.01, ***p<0.001, ****p<0.000. **E**, Univariate Cox-proportional hazard regression analysis (corrected for molecular subtype) for each Cm significantly associated with survival from the Bueno cohort. **F**, Common PM CNV (right) interaction with cancer programs (bottom) as computed by median of per-sample Spearman correlations between each Cm and CNV score (Methods). Left bar shows median of per-sample Spearman correlation to the scS-score. **G,** Distribution of samples from the Bueno cohort scored by the scS-score and grouped by FISH staining of chr22 reported as deleted or normal in the Bueno cohort. *P* value was computed using Welch Two Sample *t*-test. **p<0.01.

The ability of our scRNA-seq analysis to separate malignant cells from other TME cell types enabled us to dissect intra-tumoral heterogeneity and cancer-intrinsic expression programs at a much higher resolution and accuracy than was previously possible in bulk studies. We used consensus non-negative matrix factorization (25) (cNMF) to identify 20 unique cancer modules (Cm1-Cm20) after careful annotation of their biological pathways based on co-expression patterns across cells and enrichment of top markers in canonical cancer expression programs (Figure S2B-D; Table S5; Methods). For example, we identified a cancer cell module (Cm17) that was predominantly expressed in sarcomatoid histology tumors and was highly similar to the bulk RNA-seq derived S score from (14) (Figure S2E). Cm17 included known sarcomatoid-associated genes (e.g., *AXL, HAPLN*, *VIM*; Table S5) as well as novel ones such as *S100A3, IGFBP6*, and *CAVIN3* that have been implicated in pancreatic, breast, and lung cancer progression, respectively (26–28). To quantify the sarcomatoid content for each sample, we scored all malignant cells for Cm17 (referred to as single-cell Sarcomatoid score or scS-score hereafter) and classified tumor samples as scS-high or scS-low based on their mean scS-score ranking (Figure 2B). To investigate the relationship between different cancer modules and scS-score, we correlated Cm scores across malignant cells and samples (Figure 2C, S2C). Malignant programs most correlated with scS-score were involved in hypoxia (Cm8; *TGFBI, VEGFA*), BMP2-driven targets (Cm11; *HPGD, SYT1*), epithelial to mesenchymal transition (EMT) (Cm19; *COL1A1, MMP2*), cell migration (Cm9; *BARX1, PODXL*), cell proliferation (Cm16; *PCNA, MKI67*) and a mixed program enriched in several pathways including EMT, glycolysis, and hypoxia (Cm14; *TGFB1*, *LOX;* Figure S2D). In contrast, malignant programs anti-correlated with the scS-score were mostly enriched in epithelioid markers (Cm2; *MSLN, ITLN1),* cell projections (Cm10; *TEAD1, WWC1)* and mesothelium markers (Cm15; *HP*, *UPK3B*). We also defined other interesting malignant programs related to immune pathways that did not show strong association with scS-score, including TNF-driven inflammation (Cm7; *NFKBIA, ATF3*), interferon response (Cm18; *ISG20, IFIT1*) and antigen-presenting (Cm20; *HLA-DR*, *HLA-DQ*). Comparing each module’s expression in sarcomatoid versus epithelioid samples in bulk cohorts showed mostly consistent trends, validating our approach (Figure 2D, S2F), highlighting how our discovery cohort can be leveraged to uncover novel cancer programs at single-cell resolution and validate their association with molecular subtypes in larger PM cohorts.

To assess if the *de novo* discovered cancer programs were associated with different disease outcomes, we performed survival analysis using both the Cox proportional hazards regression analysis (adjusted for molecular subtype or histology) and the Kaplan-Meier model within each histology. We found that sarcomatoid Cm17, cell proliferation Cm16, cell migration Cm9, and mixed program Cm14 were predictive of poor outcome in both validation cohorts (Figure 2E, S2G-H). When stratified by molecular subtype we also found PTEN-signaling Cm5 and chromatin organization Cm12 to be prognostic of lower overall survival only in epithelioid and sarcomatoid Bueno cohort patients, respectively (Figure S2I).

Lastly, we performed a new computational analysis that systematically uncovers genomic interactions between Cms and expression of genes in frequently deleted PM CNV domains (Figure 2F; Methods). For instance, chromosome 22 (chr22) was inversely correlated with expression of several Cms including the epithelioid Cm2 and was positively correlated with the scS-score. Interestingly, we observe a similar trend in the Bueno bulk cohort at both expression (Figure S2J) and DNA level, as quantified by fluorescence in situ hybridization (FISH; Figure 2G), suggesting that chr22 deletions may occur preferentially in low scS-score, epithelioid-like PM tumors.

In summary, single-cell dissection of malignant cell heterogeneity uncovered genomic alterations and cancer-intrinsic gene expression programs in PM associated with different molecular subtypes, including a sarcomatoid, cell proliferation, cell migration, and mixed programs associated with poor outcome.

### Fetal-like, scS-score associated endothelial cells likely contribute to angiogenesis

The scRNA-seq data also presented the opportunity to characterize the stromal cell subsets and interactions across our cohort, which has been largely understudied in PM compared to the malignant and immune cell compartments. Based on Louvain clustering and expression of canonical markers we identified six mesenchymal and endothelial cell (EC) subsets: artery, *PLVAP*+ EC, vein, lymphatic EC (LECs), cancer-associated fibroblasts (CAFs), and smooth muscle cells (SMCs) (Figure 3A-B, S3A). Using cNMF, we identified 6 EC gene modules (Ems), where only Em3, *PLVAP*+ EC module was correlated with the cancer-intrinsic scS-score (Figure S3B-C). cNMF also detected 6 CAF modules (Fms): *COL6A2*^high^ *PNISR*^high^ (Fm1), *IGFBP6*^high^*MFAP5*^high^ (Fm2), *CDH2* ^high^*FABP5* ^high^ (Fm3), *COL16A1*^high^*COL8A1*^high^ (Fm4), *TXNIP* ^high^ *SERPING1*^high^ (Fm5), and *IGFBP2*^high^ (Fm6) (Figure S3D-E). Comparison with mesenchymal cells from normal lung scRNA-seq data (29) revealed that Fm1 and Fm2 were most similar to the adventitial fibroblasts, Fm4 to the alveolar fibroblast, Fm3 to pericytes, and Fm5 and Fm6 to Lipofibroblasts (Figure S3F).

**Figure 3.**
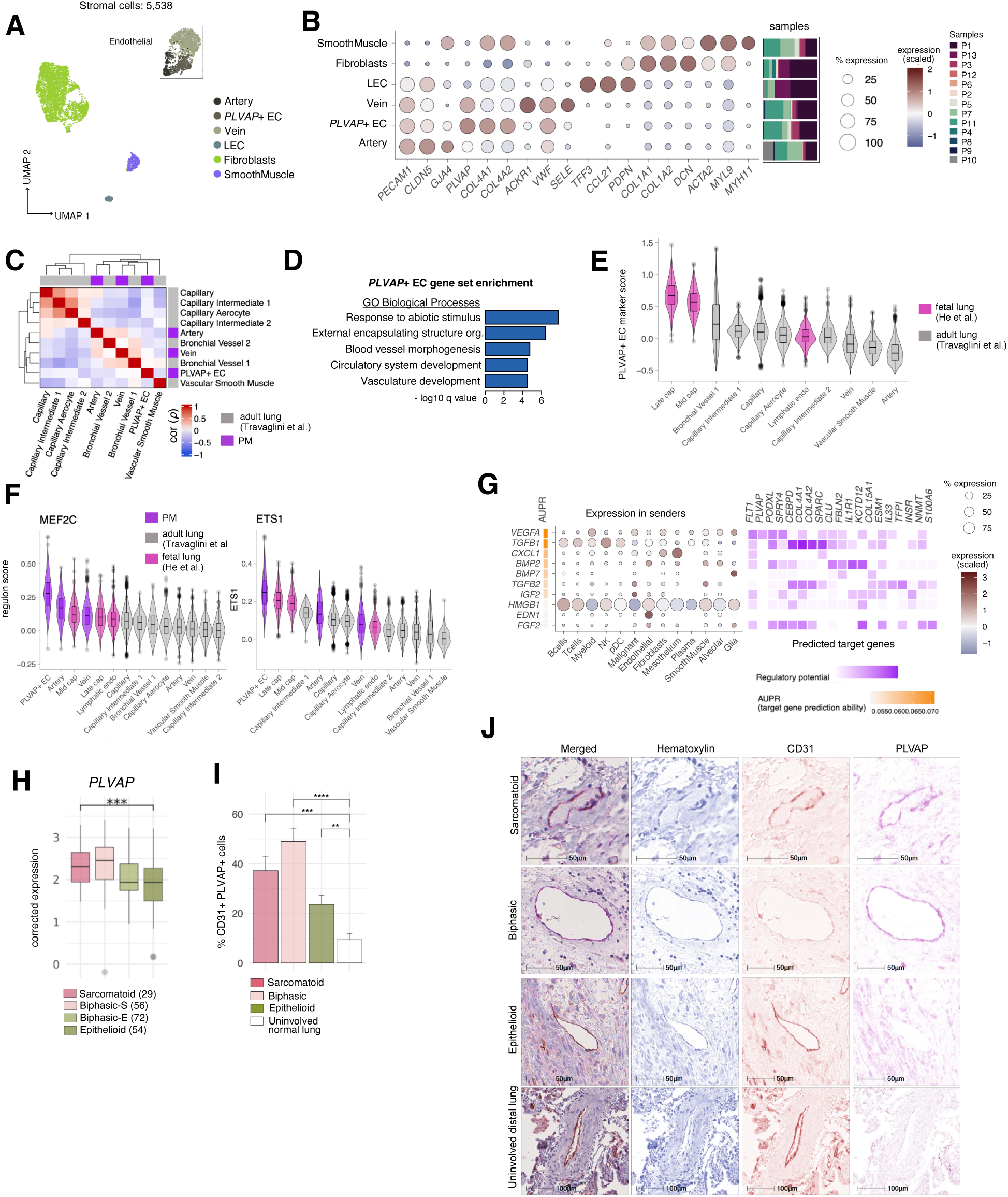
Fetal-like, cancer-enriched *PLVAP*+ endothelial cells associate with angiogenesis. **A**, UMAP embeddings of PM stromal and endothelial cell types integrated across patients. **B**, Dot plot showing expression and percentage of cells expressing top selected markers per cell type annotation and relative sample composition (right, stacked bar plots). **C**, Spearman correlation coefficient heatmap clustering the average expression profiles of endothelial cell subsets found in normal adult distal lung and PM samples. **D**, Gene set enrichment analysis of *PLVAP*+ EC markers compared against the Gene Ontology (GO) biological processes database. Top five enriched categories were displayed. **E**, Distribution of *PLVAP*+ EC marker score in fetal and adult distal lung endothelial cell subsets, ordered from highest to lowest median score. **F**, Distributions of the *MEF2C* (left) and *ETS1* (right) regulon activity in fetal adult distal lung, and PM endothelial cell subsets. **G**, NicheNet prediction of ligand prioritization (top 10 displayed), their abundance in sender cell types (left dotplot), and their cognate targets among *PLVAP*+ EC markers (right heatmap). **H**, Sample distributions of *PLVAP* expression in the Bueno cohort grouped by molecular subtype after correcting for endothelial content. *P* values were computed comparing sarcomatoid and epithelioid subtypes using Welch Two Sample *t*-test. p<0.001 = ***; **I**, Quantification of IHC staining of PVLAP+ CD31+ endothelial cells in PM tumor tissue sections of sarcomatoid (n=2), biphasic (n=2), and epithelioid histology (n=2) compared with normal adjacent distal lungs (n=4). Between 9 and 23 regions of interest (ROIs) were quantified for each sample. *P* values were computed comparing each PM subtype to the normal tissue using Welch Two Sample *t*-test. *p<0.05, **p<0.01, ***p<0.001, ****p<0.0001. **J,** Representative micrographs from tissue sections from patients with sarcomatoid, biphasic, and epithelioid PM histologies and uninvolved normal distal lung tissue section stained with anti-PLVAP (purple) CD31(brown) and hematoxilyn (blue).

Integration with normal lung EC scRNA-seq data (29) similarly confirmed high correspondence between normal and PM EC subsets, except for the *PLVAP*+ EC population (Figure 3C). To examine the functional role of this EC subset we performed gene set enrichment analysis and found high enrichment of genes associated with blood vessel morphogenesis and development (Figure 3D). This suggested that *PLVAP*+ ECs may be more prominent in development and prompted us to compare this population to a recently published fetal lung single-cell atlas (30); indeed, top markers expressed in *PLVAP*+ ECs were also highly expressed in distal fetal lung endothelial populations relative to EC subset from adult lungs (29) (Figure 3E). *PLVAP* was recently reported as a marker for fetal-like ECs in hepatocellular carcinoma, but other marker genes (e.g. *COL4A1/2*, *RGCC*, *HSPG2*, *COL15A1*) were unique to this population arguing that this is an PM-specific, fetal-like EC subset (31).

Next, we employed single cell regulatory network and clustering (SCENIC) (32) to decipher the key transcription factors (TFs) and downstream gene regulatory modules (regulons) for each EC subset (Figure S3G). This analysis revealed *ETS1* and *MEF2C* to be among the top TF regulators of *PLVAP*+ EC population, including 318 and 63 genes in their regulons, respectively. Both *de novo* identified *ETS1* and *MEF2C* regulons were most highly expressed in the *PLVAP*+ EC and fetal EC subpopulations (Figure 3F; Table S6). In agreement, *ETS1* and *MEF2C* are known to be required for endothelial patterning in embryonic angiogenesis and VEGF-stimulated EC migration in mouse models (33–35).

Given that *ETS1* and *MEF2C* are also known to regulate angiogenesis (36,37), we speculated that *PLVAP*+ ECs may play an important role in angiogenesis downstream of VEGF signaling. To examine which TME signaling pathways are most likely to regulate the gene expression of *PLVAP*+ ECs we employed NicheNet (38) and found that *VEGFA* was indeed the top predicted ligand, expressed predominantly in PM myeloid and tumor cells (Figure 3G). Not surprisingly, *PLVAP*+ ECs also showed the highest expression of VEGFA receptors *KDR* and *FLT4* (Figure S3H). Worth noting, in the Hmeljak validation cohort the combined expression of highly specific markers for *PLVAP+* EC subset was significantly correlated with poor survival (Figure S3I).

Furthermore, *PLVAP*+ EC expression was enriched in scS-score high, non-epithelioid PM in bulk RNA cohorts after correcting for endothelial content (Figure 3H, Figure S3J). To experimentally validate the presence of *PLVAP*+ ECs in PM and enrichment in non-epithelioid tumors, we performed dual immunohistochemistry staining for PLVAP/CD31 on tissue sections derived from PM patients encompassing all three histological subtypes, along with uninvolved normal distal lung tissue (control) obtained from patients with lung adenocarcinoma. The quantified percentages of endothelial cells exhibiting concurrent expression of CD31 and PLVAP within blood vessels were significantly increased in PM compared to control tissue, with the largest differences observed in non-epithelioid PM (Figure 3I-J). Taken together, we discovered a PM-specific, fetal-like, angiogenic *PLVAP*+ EC subset that is likely regulated by TF *ETS1*, *MEF2C*, and *VEGFA* signaling; this population specifically expresses VEGFA receptors *KDR* and *FLT4* and is enriched in non-epithelioid PM tumors, which likely favors tumor survival and contributes to a worse disease outcome. These findings thus support further investigation of anti-VEGFA agents (7) in patients with PM with high *PLVAP*+ EC abundance.

### Macrophages in scS-high PM express *CXCL9/10/11* and likely contribute to T-cell infiltration

To characterize the diversity of myeloid cells in PM, we performed unsupervised clustering followed by integration and annotation of cell subtypes based on canonical markers (Figures 4A-B, S4A-C). We identified 8 different myeloid subsets such as dendritic cells, further separated into cDC1, cDC2 and mregDCs, plasmacytoid dendritic cells (pDCs), classical (CD14+) and non-classical (CD16+) monocytes, mast cells, and a large and heterogeneous cluster of tumor-associated macrophages (TAMs). We observed that *VISTA*, an immune checkpoint (IC) gene shown to be preferentially expressed in epithelioid subtypes (10,39) was most highly expressed by monocytes amongst myeloid subsets and all other cell types and higher in CD14+ monocytes in scS-low epithelioid tumors (Figure 4C). *VISTA* has been targeted in clinical trials for PM and quantifying its expression at a single-cell resolution can provide insight into its potential therapeutic mechanisms and how these differ across histological subtypes.

**Figure 4.**
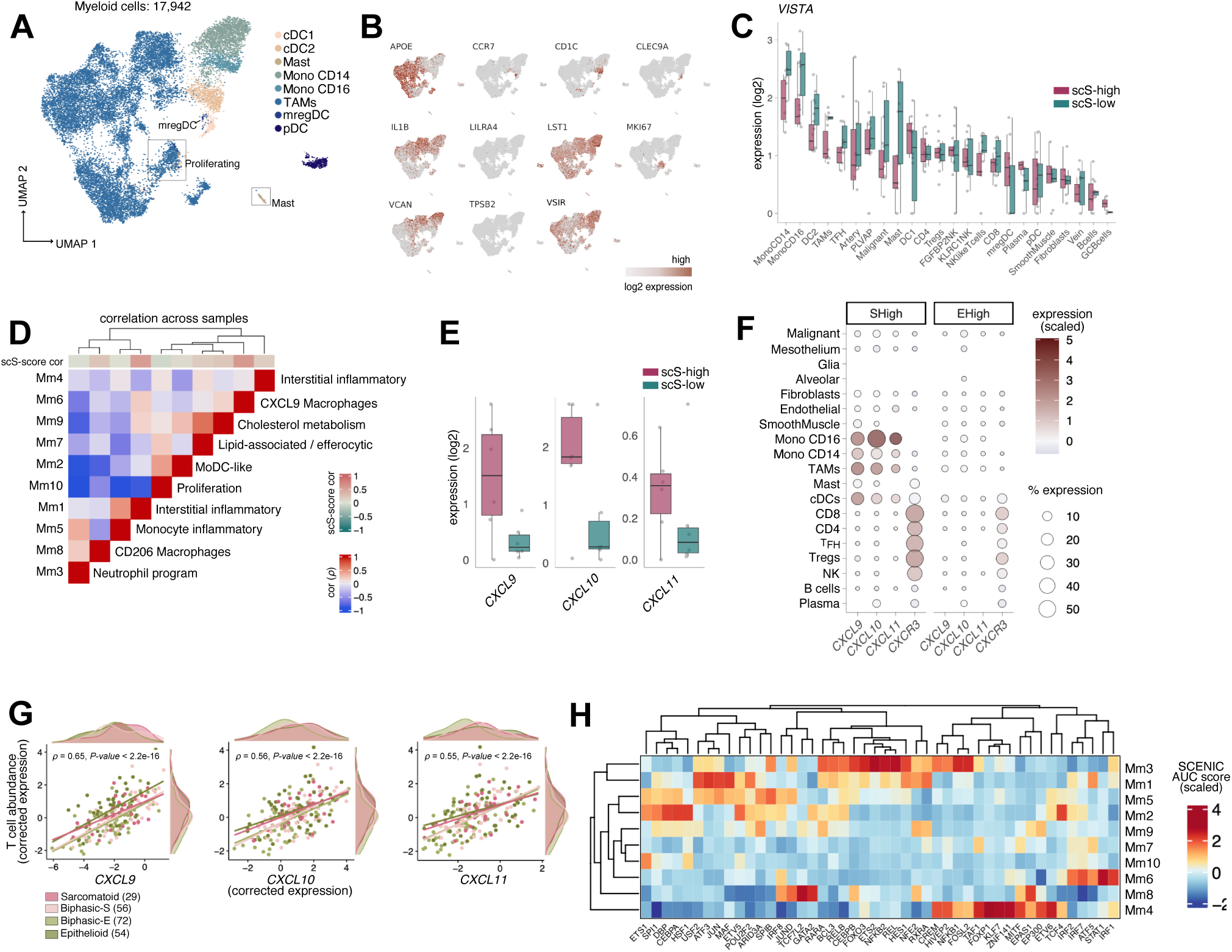
Macrophages in scS-high PM express *CXCL9/10/11* and likely contribute to T-cell infiltration. **A**, UMAP embeddings of PM myeloid cells integrated across patients. **B**, Feature plots of key markers used for myeloid cell type annotation. **C**, Sample distributions of log2-normalized expression levels of *VSIR (VISTA)* across all PM cell types, including myeloid subsets and split by scS-high and scS-low groups. FDR-adjusted *P* values were computed using Welch Two Sample *t*-test. **D**, Pairwise Spearman correlation of sample-averaged scores derived from the 10 Mms identified in PM TAMs. Each Mm was annotated with the most representative biological pathway. Top color bar shows correlation of each Mm to the scS-score. **E**, Sample distributions of log2-normalized mean TAM expression of *CXCL9/10/11* split by scS-high and scS-low samples. FDR-adjusted *P* values were computed using Welch Two Sample *t*-test. **F** Dot plot showing expression and percentage of cells expressing *CXCL9/10/11* and their receptor *CXCR3* across cell types and split by scS-high and scS-low samples. **G**, Expression of *CXCL9/10/11* versus T cell abundance inferred as the average expression of T cell marker genes in the Bueno cohort corrected for immune content. Spearman correlation *P* values are shown. **H**, Heatmap of the SCENIC significant regulon activities (scaled AUC score) and correspondent TFs (columns) in each TAM subset. TAM = tumor-associated macrophage; Treg = regulatory T cell; T_FH_ = T follicular helper cell. *p<0.05, **p<0.01, ***p<0.001, ****p<0.0001.

When applying cNMF to dissect TAM heterogeneity we detected 10 macrophage modules (Mm), including an interstitial macrophage-like state (Mm1; *SELENOP, LYVE1*), an inflammatory *CXCL9^high^* TAM state (Mm6; *C1QC, STAT1*), and lipid-associated TREM2^high^ TAM state (Mm7 and Mm9; *GPNMB, SPP1, HILPDA, TREM2*) (Figure 4D, S4D-E). We find that Mm1 and Mm6 were most correlated with the scS-score (Figures 4D) and, in agreement, *CXCL9/10/11* expression was higher in scS-high versus scS-low myeloid cells (Figure 4E-F). These chemokines are known to bind receptor CXCR3, recruit T cells to the tumor core, and correlate in expression with lymphocyte abundance in melanoma and lung cancer (40,41). In our cohort *CXCL9/10/11* were most highly expressed in monocytes and TAMs, while their corresponding receptor *CXCR3* was specifically expressed NK and T cells, especially in CD8 and regulatory T (Treg) cells (Figure 4F). Increased recruitment of T cells in scS-high PM tumors via these interactions is further supported by significant correlations between *CXCL9/10/11* expression and T cell abundance in the Bueno cohort (Figure 4G).

To further investigate the regulation underlying different myeloid and TAM subsets, we performed regulon analysis using SCENIC (Figures 4H, S4F). This *de novo* analysis captures the known role of *IRF7* in pDC function (42) and predicts regulons driven by TFs *MAFB, MEF2C, BCLAF1* and *YY1* in monocytes (Figure S4F). Additionally, TFs *MAF, ATF3*, and *JUN* were enriched for regulon activity with scS-high associated Mm1 TAM state, while known IFNγ signaling TFs *STAT1* and *IRF1* (43) were predicted as regulators of Mm6 and *CXCL9/10/11* expression (Figure 4H).

In summary, we observe differences in myeloid expression associated with different PM subtypes, including higher *VISTA* expression in scS-low monocytes and increased production of *CXCL9/10/11* chemokines implicated in chemotaxis of T cells in scS-high TAMs likely regulated by TFs *STAT1* and *IRF1*; these findings can inform on future immunomodulatory therapies targeting myeloid cells in PM.

### Molecular dissection of T cell programs and IC molecules shows association with scS-score

To comprehensively characterize the T and NK cellular diversity in PM *de novo*, we again utilized two complementary unsupervised clustering approaches—Louvain clustering and cNMF. Louvain clustering identified major cell subsets in the tumor samples including CD4, CD8, Treg, T follicular helper (T_FH_) cells, and two NK cell subsets marked by high expression of *KLRC1* and *FGFBP2* (Figures 5A-B, S5A). Using cNMF we additionally uncovered functional T cell expression modules (Tms), such as, naïve (Tm1), stress response (Tm8), interferon response (Tm12), inflammatory (Tm3), gamma delta (Tm9), and proliferative T cells (Tm10; Figures 5C, S5B-C). We found five T cell modules positively correlated with scS-score, including a Treg-associated program (Tm7; *FOXP3* and *IL2RA*) and four other modules linked to CD8 cell states—namely, progenitor (Tm11; *XCL1, GNG4*), exhaustion (Tm5; *HAVCR2, LAG3*), effector (Tm2; *NKG7, GZMA*), and MHC II genes expressing module (Tm4) linked to CD8 T cell activation (44) (Figure 5C-D). Effector, exhaustion, and Treg modules showed increased expression in T cells from scS-high tumors and were significantly enriched in bulk deconvolution analysis comparing sarcomatoid versus epithelioid tumors after correcting for T cell content and selecting specific markers in the validation cohorts, arguing that higher immune infiltration in scS-high tumors is accompanied by a shift toward CD8 and Treg fractions (Figure 5E, S5D). Increased exhaustion in scS-high T cells was also supported by higher expression of *HAVCR2* and *LAG3* as well as known IC targets, including *PDCD1*, *TIGIT*, and *CTLA4* (Figure 5F), as previously reported in bulk RNA-seq studies (13,14). Amongst T cell subsets, *CTLA4* and *TIGIT* showed highest expression in Tregs, while *PDCD1* was most highly expressed in CD8 and T_FH_ cells in scS-high and scS-low tumors, respectively (Figures 5G, S5E).

**Figure 5.**
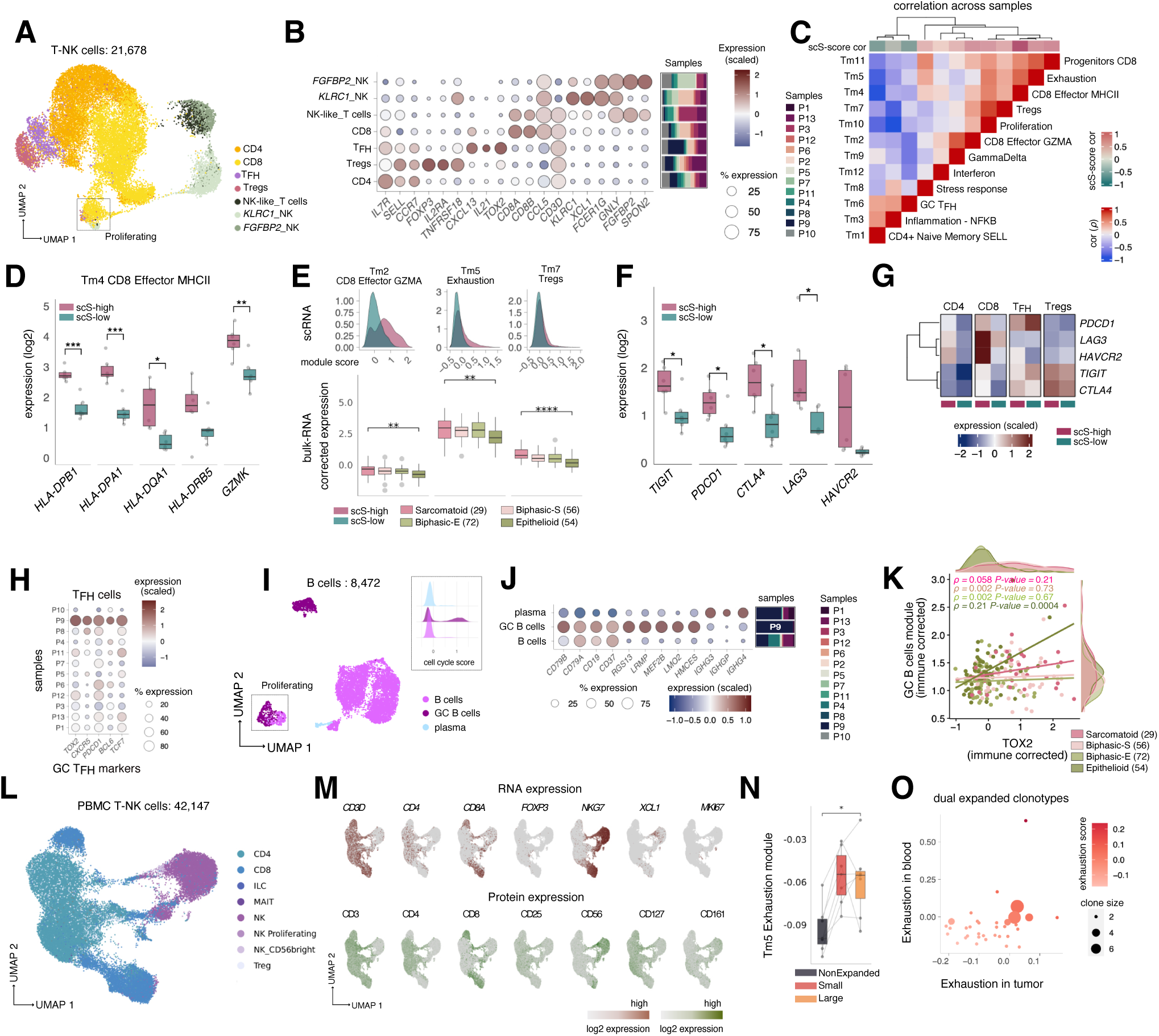
Molecular dissection of T cell programs and IC molecules shows association with scS-score. **A**, UMAP embeddings of PM tumor T and NK cells integrated across patients. **B**, Dot plot showing the expression and percentage of cells expressing key markers used for cell type annotation with relative sample composition for each cell type (right, stacked bar plots). **C**, Pairwise Spearman correlation of sample-averaged scores derived from the 10 Tms identified in T cells. Each Tm was annotated with the most representative biological pathway. Top color bar shows correlation of each Tm to the scS-score. **D** Sample distributions of log2-normalized mean T cell expression of selected genes from Tm4 averaged per samples and split by scS-high and scS-low samples. FDR-adjusted *P* values were computed using Welch Two Sample *t*-test. **E,** Cell distributions of Tm module scores in all scS-high and scS-low samples (top) and sample distributions of Tms expression in the Bueno cohort for each PM molecular subtype after correcting for immune content (bottom). FDR-adjusted *P* values were computed comparing sarcomatoid and epithelioid subtypes using Welch Two Sample *t*-test. **F**, Sample distributions of log2-normalized mean T cell expression of known IC molecules. FDR-adjusted *P* values were computed using Welch Two Sample *t*-test. **G**, Scaled log2-normalized mean expression of known IC molecules across T cell subsets in scS-high and scS-low samples. **H,** Dot plot showing the expression and percentage of cells expressing markers for germinal center T follicular helper cells identified in Tm6. **I,** UMAP of B-cells compartment, including germinal center B cells (GC-B cells) predominantly found in P9. Inset shows distribution of cell cycle scores grouped by cell type annotations. **J,** Dot plot showing the expression and percentage of cells expressing top markers of B cells subsets and relative sample composition (right, stacked bar plots). **K,** Average expression of marker genes of GC B cells (y-axis) versus expression of TOX2 (identifying GC T_FH_ cells, x-axis) in the Bueno cohort corrected for immune content. Spearman correlations and relative *P* values were computed for each molecular subtype. **L,** UMAP embedding of annotated T and NK cell subsets integrated across all PBMC samples. **M,** Feature plots of T and NK cell subsets representative markers (top, RNA; bottom, protein). **N,** Sample distribution of the mean exhaustion score in non-expanded vs expanded clonotype CD8 cells from PBMC samples (Methods). *P* value was computed using Welch Two Sample *t*-test. **O,** Dual expanded clonotypes as identified both in patient-matched tumor and PBMC samples distributed by their exhaustion scores in the corresponding sample source. *p<0.05, **p<0.01, ***p<0.001, ****p<0.0001.

We observed expression of germinal center (GC) T_FH_ cell markers (e.g., *TOX2*, *CXCR5*) (45) in sample P9 enriched module Tm6 (Figure 5H), prompting us to examine the B cell compartment where we also identified a population of highly proliferating GC B cells found almost exclusively in P9 (Figure 5I-J). Notably, enrichment of both GC T_FH_ and B cells suggests the presence of mature tertiary lymphoid structures (TLS) in this epithelioid PM patient. To investigate a link between TLS presence and molecular subtypes, we correlated expression of GC T_FH_ marker *TOX2* with the top markers of GC B cells (Figure 5J) and found significant association only in epithelioid samples (Figure 5K; *r =*0.46, *P* value *=* 0.0004). Interestingly, a previous study showed histological evidence of TLS presence in a subset of epithelioid PM tumors associated with longer survival (46).

Next, we examined the CITE-seq and scTCR-seq data for all PBMC lymphocytes, which showed consistent RNA and protein expression (Figure 5L-M, S5F) and detection of more than 3,000 expanded clonotypes in CD8 T cells (Figure S5G). Expanded TCR clonotypes may be indicative of reactive CD8 T-cells recognizing tumor antigens or bystanders CD8 memory T cells but only the former may lead to terminal exhaustion (47). Hence, we scored CD8 T cells with detectable TCR sequence by the exhaustion module previously identified in tumor-infiltrating lymphocytes (Tm5) and found that expanded CD8 clonotypes have significantly higher exhaustion score compared to non-expanded clonotypes (Figure 5N). We also show significant increases for activation (Tm4) and cytotoxicity (Tm2) CD8 module scores (Figure S5H). These trends, albeit not significant perhaps due to smaller sample size, were also observed in tumor infiltrated T cells, where expanded clonotypes also mapped primarily to CD8 T cells and made up a higher fraction of CD8 T cells in scS-high tumors (Figure S5I-K). Finally, we identified several expanded clonotypes present in both tumor and blood patient-matched samples and exhibiting high exhaustion scores, further suggesting systemic anti-tumoral T cell activity (Figure 5O, S5L-M).

Taken together, molecular characterization of B and T cells revealed a higher abundance of Tregs, expression of IC targets, and CD 8 exhaustion, cytotoxicity and activation modules associated with the scS-score; in contrast, germinal center T_FH_ and B cells markers suggest preferential TLS formation in epithelioid PM tumors.

### NK cell IC blockade targeting NKG2A as a novel therapeutic strategy in PM

In the past decade, immunomodulatory drugs have become a mainstay for the treatment of cancer (48), including anti-PD1 and anti-CTLA4 combination therapy recently approved for use in PM (49). NK cells have been largely unexplored in PM but also represent a viable therapeutic target (50). We found a significant survival benefit of higher NK cell infiltration in tumors from patients with epithelioid PM (Figure 6A, S6A) and observe a similar trend when using a Cox proportional hazard regression model across all subtypes in both validation cohorts (Figures S6B-C). Combined with our previous observation of higher NK cell infiltration in epithelioid PM (Figure 1E), this analysis indicates that NK cell abundance may represent an important, epithelioid-specific prognostic biomarker. To dissect the crosstalk between NK and malignant cells and identify new therapeutic avenues, we curated a list of NK cell inhibitory receptors and cognate ligands and found that *KLRC1* and its ligand *HLA-E* were both highly expressed by NK and malignant cells, respectively, in comparison to other ligand receptor pairs in our scRNA-seq data (Figure 6B). Additionally, we found that the fraction of *KLRC1*-expressing NK cells was most abundant in PM compared to other cancer types after integration with scRNA-seq data from a pan-cancer immune cell atlas (51) (Figure 6C).

**Figure 6.**
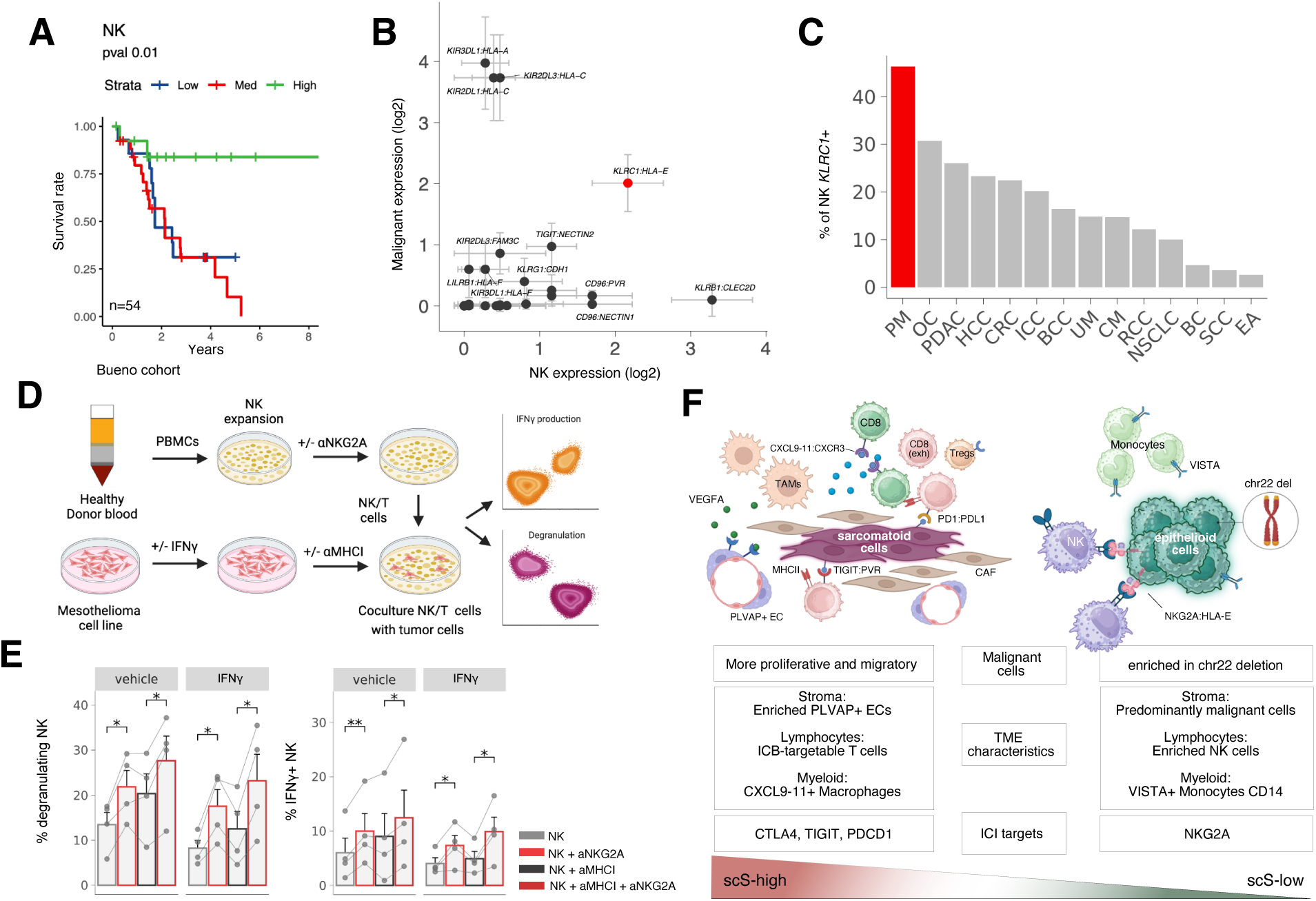
NK cell IC blockade targeting NKG2A as a novel therapeutic strategy in PM. **A**, Kaplan-Mayer curve stratifying epithelioid samples in the Bueno cohort by NK cell abundance (deconvolved using BayesPrism). **B,** Log2-normalized mean expression of ligand-receptors expressed by malignant cells (y axis) and NK cells (x axis) respectively. Error bars represent standard errors of expression across samples. **C**, Percentage of *KLRC1* expressing NK cells averaged across patients with different cancer types combining scRNA-seq data from our and a pan-cancer study (51). **D,** Schematic of the co-culture experimental design. **E,** Activation of NK cells co-cultured with mesothelioma cell lines upon NKG2A blockade, indicated by degranulation (left) and IFNγ production (right) with or without anti-MHCI antibody and with or without IFNγ stimulation of PM cell lines. *P* values were computed using a paired Student’s *t*-test. Error bars represent standard error. **F**, Schematic of the key TME differences between scS-high and scS-low PM elucidated in our study. BC = breast cancers; BCC = basal cell carcinomas; CM = cutaneous melanoma; CRC = colorectal cancers; EA = endometrial adenocarcinomas; HCC = hepatocellular carcinomas; ICC = intrahepatic cholangio carcinomas; PM = pleural mesothelioma; NSCLC = non-small-cell lung cancers; OC = ovarian cancers; PDAC = pancreatic ductal adenocarcinomas; RCC = renal cell carcinomas; SCC = oropharyngeal squamous cell carcinomas; UM = uveal melanoma. *p<0.05, **p<0.01, ***p<0.001, ****p<0.0001.

Antibodies (monalizumab) targeting NKG2A (encoded by *KLRC1* gene) has been shown to enhance both NK and CD8 T cell response (50). To experimentally test if blocking NKG2A-HLA-E interaction could augment NK cells anti-tumor function in PM, we co-cultured four mesothelioma cell lines with blood-derived NK cells in the presence or absence of anti-NKG2A antibody (Figure 6D, S6D). Flow cytometry analysis showed that the mesothelioma cell lines constitutively express HLA-E, which increased following IFNγ treatment (Figure S6E), while the NK cells expressed high levels of NKG2A (Figure S6F). Next, NK cells were co-cultured with the mesothelioma cell lines for 16 hours in the presence or absence of anti-NKG2A antibody, and IFNγ production and degranulation (CD107a^+^, Granzyme A^-/low^) were measured thereafter by flow cytometry as read-outs for NK cell activation. We found that NKG2A blockade significantly increased NK degranulation and IFNγ production, regardless of whether the tumor cell lines were pre-stimulated with IFNγ to increase HLA-E expression (Figure 6E). These differences remained significant after applying a boolean operator for gating on total activated NK cells undergoing degranulation or producing IFNγ (Figure S6G). We tested this interaction also in the presence of anti-MHC class I (MHCI) antibody since expression of MHCI on tumor cells is known to suppress NK cell activation. This additional step confirmed that enhanced NK cell activation was indeed primarily due to the targeted blockade of the NKG2A-HLA-E interaction (Figures 6E, S6G).

In conclusion, our analysis demonstrates that NK cell infiltration is a prognostically relevant biomarker in epithelioid PM subtypes, and that targeting NKG2A significantly augments NK cells tumor cytotoxicity, warranting further investigations as a viable immunotherapy strategy in PM.

## DISCUSSION

We performed scRNA-seq profiling of ~140,000 human tumor and peripheral blood cells and identified 54 gene expression modules across cellular compartments to generate the first single-cell sequencing atlas of PM. Analysis of malignant cell heterogeneity showed presence of all 4 molecular subtypes in biphasic and most epithelioid PM tumors, supporting the notion that PM tumors do not classify into discrete molecular subtypes but rather lie on a continuum between sarcomatoid and epithelioid histology (13,14). Consequently, we adopted a rank-based analytical strategy designed to capture pairwise enrichment of different cellular programs across patients, which uncovered a highly distinct TME associated with a single-cell resolution, cancer-intrinsic sarcomatoid signature, we termed scS-score (Figure 6F). We also uncovered cell migration, proliferation, and mixed hypoxia/EMT cancer modules that were associated with high scS-score across patients and predictive of poor outcome. In contrast, cancer modules containing epithelioid markers were associated with chromosome 22 deletion in our scRNA-seq data, which was supported by RNA expression and DNA FISH data from the Bueno validation cohort. A multi-regional whole exome-sequencing from 22 PM patients aiming at reconstructing clonal trajectories reported chr22 deletion as a late event during PM evolution but did not find an association with the epithelioid subtype perhaps due to the small sample size (52), whereas our predictions leveraged on CNV-gene module co-variation analysis across malignant cells.

Our *de novo* analysis led to the discovery of a fetal-like, *PLVAP*+ endothelial cell population, which we predict to be responsive to VEGFA signaling through receptors KDR and FLT4 and promote angiogenesis. This population was enriched in PM tumors when compared to ECs from normal adult lungs and was also associated high scS-score (scS-high) samples, which we validated by IHC. Bevacizumab, a monoclonal antibody targeting VEGFA effective in the treatment of many cancers (53), has been introduced in first-line standard of care for patients with unresectable PM albeit with limited benefits (7). Efforts in identifying biomarkers of treatment response have focused on plasma levels of VEGF-A (pVEGFA) and molecules eliciting similar angiogenic responses with inconclusive results (54,55). It is tempting to speculate that this population of *PLVAP*+ ECs may represent a novel biomarker for anti-angiogenic therapy response and a putative future drug target to abrogate tumor-induced angiogenesis.

Examination of the immune composition of PM samples with high scS-score showed a higher proportion of Tregs and CD8 effector and exhausted T cells, in line with past bulk-RNA studies (12,13), and further uncovered a population of CD8 MHC II^+^ T cells, which was previously reported to induce pro-inflammatory activity in patients responding to neoadjuvant chemotherapy in breast cancer (44).We also provide molecular evidence for T_FH_ cells positive for *CXCL13* and *IL21,* which are relevant biomarkers of immunotherapy response (56), and further describe a patient-specific *TOX2*+ T_FH_ transcriptional program associated with the presence of highly-proliferating germinal center B cells that could signify the presence of mature TLSs (52). Indeed, *TOX2* has been shown to be essential for maintaining a T_FH_ phenotype in *ex vivo* GC T_FH_ isolated from human tonsils (45). Supporting our finding, TLSs have been previously observed in PM using bulk-RNA and histological analysis on a cohort of 123 chemo-naive patients, which was linked to improved survival and enriched in epithelioid tumors (57). Finally, we identified a *CXCL9/10/11* expressing TAM population in PM that is associated with high scS-score samples and likely contributes to chemotaxis for T cell trafficking to the tumor core. Consistent with this observation, *CXCR3* (receptor for *CXCL9/10/11*) was more expressed in scS-high tumor T cells, especially in CD8 T cells and Tregs that have higher abundance in scS-high tumors. Further supporting this finding is a recent study employing spatial transcriptomics in PM biphasic samples, which showed increased lymphocytic infiltrate and expression of chemokines *CXCL9/10* in sarcomatoid-enriched regions (58). Future time-course studies will be needed to decipher the precise molecular events that trigger these highly divergent TMEs that track with the sarcomatoid-epithelioid axis.

Resolving the complexity of the immune-stroma-tumor interface and composition in the TME is of high clinical significance given that there are over 4700 immunotherapy agents in development (48), emphasizing the need for rational clinical trial design and patient treatment stratifications based on observations such as those reported here. Our data-driven approach highlighted an immunosuppressive NKG2A-HLA-E interaction between NK and tumor cells, which enhanced NK cell degranulation and IFNγ production upon NKG2A blockade in co-cultures with 4 PM cell lines. A previous study similarly reported reactivity of NK cells isolated from PBMC of healthy individuals against mesothelioma cell lines when stimulated with IL-15 (59). Further supported by the findings that *KLRC1* expression in NK cells is more abundant in PM relative to other cancer types and that NK cell content is an indicator of better overall survival in epithelioid PM, this initial finding lays the ground for further investigations in experimental models of PM using anti-NKG2A therapeutics (e.g., monalizumab).

In conclusion, this study demonstrates the potential of high-throughput cellular profiling via scRNA-seq and in-depth analysis on PM clinical samples in identifying new cellular programs, prognostic signatures of disease outcome, and therapeutic targets towards the goal of achieving more effective, personalized therapies in PM.

### Study limitations

Our study comes with several limitations. Firstly, the small sample size of this rare pleural cancer limited our ability to sample patients evenly across different molecular subtypes. Our analysis strived to overcome this limitation by corroborating our main findings using large bulk RNA-seq cohorts and performing associations between cell composition and gene expression programs using rank statistics (Spearman correlation), which takes advantage of the power of the entire cohort rather than dividing samples into discrete groups. Secondly, differences in TME along the sarcomatoid to epithelioid subtype gradient in our study were investigated using gene expression information alone, whereas a recent study employing bulk multimodal molecular profiling reported orthogonal axes of molecular divergence driven by DNA methylation, genomic ploidy, and immune infiltration (12). Future efforts in characterizing PM should aim to leverage such multimodal technologies at a single-cell resolution. Thirdly, even though we were able to identify and validate the presence of a fetal-like, endothelial subpopulation, the stromal component in our scRNA-seq cohort was overall underrepresented, accounting for only 6,352 cells with several samples having very low numbers. This may have precluded us from uncovering additional stromal subtypes of relevance for PM progression, especially dissection of cancer-associated fibroblasts (CAF) that are known to be abundant in PM and contribute to its pathogenesis (60). Fourthly, we capture 3,214 NK cells in our data that form 3 distinct populations; however, we anticipate that higher sampling of these cell types in PM and across cancer types will better inform on their functional diversity and therapeutic potential (61). Lastly, our single-cell catalogue does not capture neutrophils, which are known to escape detection in human samples utilizing the 10X Chromium scRNA-seq platform.

## METHODS

### Human tumor sample collection

Tumor samples were obtained from diagnostic biopsies and surgical specimens of patients undergoing resection at Mount Sinai Hospital after obtaining informed consent in accordance with a protocol reviewed and approved by the Institutional Review Board at the Icahn School of Medicine at Mount Sinai (IRB Human Subjects Electronic Research Applications 10-00472 and 10-00135) and in collaboration with the Biorepository and Department of Pathology. Clinical information of subjects can be found in Table S1. Only patients with treatment-naive PM were included in this study.

### Tumor sample processing

Tumor samples were transported in MACS® Tissue Storage Solution stored at 4°C, rinsed with PBS, minced and incubated in a rotation shaker for 40 minutes at 37°C in Collagenase IV 0.25mg/ml, Collagenase D 200U/ml and DNAse I 0.1mg.ml (all Sigma). Cell suspensions were then aspirated through a 18G needle ten times and strained through a 70-micron mesh prior to RBC lysis. Dead cells were removed using an EasySep Dead Cell Removal (Annexin V) Kit. Cell suspensions were sorted into CD45+ and CD45-cells using the EasySep™ Human CD45 Depletion Kit per kit instructions.

### Tumor single-cell library construction and sequencing

Single-cell RNA-seq (scRNA-seq) was performed on tumor samples using the Chromium platform (10x Genomics, Pleasanton, CA) utilizing both the 3′ and 5’ gene expression kits. Approximately 6000 CD45^+^ and CD45^-^ cells were loaded into each channel of the 10x Chromium controller, following the manufacturer-supplied protocol. For 5’ gene expression samples, BCR and TCR CDR3 sequences were enriched using the human V(D)J B/T cell enrichment. 10x libraries were constructed using the 10x supplied protocol and sequenced at the Mount Sinai Genomics Core Facility. Gel-bead in emulsions (GEMs) were generated on the sample chip in the Chromium controller. Barcoded cDNA was extracted from the GEMs using Post-GEM RT-cleanup and amplified for 12 cycles. Amplified cDNA was fragmented and subjected to end-repair, poly-A-tailing, adaptor ligation, and 10X-specific sample indexing following the manufacturer’s protocol. Libraries were quantified using Bioanalyzer (Agilent) and QuBit (Thermofisher) analysis and then sequenced in paired-end mode on a HiSeq 2500 instrument (Illumina, San Diego, CA).

### PBMC sample processing and sequencing

PBMCs were isolated within 3 hours of collection via Ficoll density gradient centrifugation for 10 min at 1200g room temperature. The supernatant was then spun down at 500g for 10 min at 4 degrees C and the pellet resuspended to a concentration of 10×10^6 cells/ml cold Human Serum AB (GemCell HAB and HAB + 20% DMSO in 1:1 ratio). The resulting PBMCs were stored in 2ml Cryogenic vials in liquid nitrogen. For cell isolation PBMCs were thawed, washed 2X in RPMI 2% FCS, treated with ACK lysis buffer (Lonza) to remove RBCs and briefly incubated with DAPI. 300,000 cells were then sorted on a DAPI negative gate. Cells were then stained for 30 minutes at room temperature with a panel of 138 Total-Seq-C antibodies (Biolegend, Stoeckius et al 2017) and washed 3x using the HT1000 laminar wash system (Curiox). Cells were then counted using the Cellaca MX High-throughput Automated Cell Counter as described in the manufacturer’s protocol (Nexcelom), pooled and loaded on the 10x Chromium 5’ V2 and Next GEM Chip K Kit using a superloading strategy mixing cells from the same sample across lanes. BCR and TCR CDR3 sequences were enriched using the human V(D)J B/T cell enrichment. Libraries were prepared according to manufacturer’s protocol (10x Genomics) and sequenced on a NovaSeq 6000 System using the S4 2x 150 kit (Illumina). Raw reads were aligned to the human transcriptome using a splice-aware algorithm to produce cell-by-gene count matrices. Cells were separated to their respective samples using a combination of public (Scrublet by Wolock et al. Cell Systems 2019) and Immunai algorithms.

### scRNA-seq data preprocessing, quality control, clustering, annotation, and differential expression

Pair-ended FASTQ files were mapped to the GRCh38 human transcriptome using the count function in CellRanger > v3.1. The count matrices obtained were normalized, log-transformed and scaled using the Seurat v4 package v4.4 in R. Cells with < 400 genes, < 1000 unique molecular identifier (UMI) counts or > 25% mitochondrial gene expression detected were removed from downstream analyses. Principal component analysis (PCA) and k-nearest neighbor (kNN) graphs were computed using Seurat default parameters. Based on the kNN graphs, a shared nearest neighbor (SNN) graph was constructed to cluster cells with the original Louvain algorithm as implemented in Seurat. High-level cellular compartment annotations were assigned to clusters based on expressions of known cell class markers. Doublets were identified in two ways: doublet clusters were identified with higher-than-average gene and UMI counts, as well as expressions of markers from multiple high-level cellular compartment (e.g., CD45+ and CALB2+) and manually removed from downstream analyses and removed from downstream analysis. Data integration within each cell compartment was performed using harmony v.0.1 to minimize sample-derived batch effects in aggregated visualizations. For batch effect correction across all cell compartments, scANVI model with n_layers=3 and n_latent=32 from scvi-tools v0.20.3 was used on raw counts to integrate the data across samples with default parameters when training. Differential expression analyses for *de novo* marker discovery were performed using Seurat FindMarkers function using a Wilcoxon Rank Sum test. For pairwise comparisons between subset groups we used muscat v1.12.1 package using DESeq2 method. Pathway and gene ontology analysis was carried out with clusterProfiler R package v4.6.0, using function enricher.

### Defining cell programs using consensus non-negative matrix factorization (cNMF)

We applied non-negative matrix factorization implemented in the Python package cNMF v1.3.4 to identify cellular states in each of the following cell types: malignant, endothelial, CAF, tumor-associated macrophages, and T cells. For each, we tested from 5 to 30 K with 100 replicate, and filtered outlier components with Euclidean distance > 0.3 from their nearest neighbors. Then based on the trade-off between reconstruction error and factorization stability and manual inspection of the modules we selected the most appropriate Ks. We then computed cNMF module scores by taking the top 20 genes ranked by spectra scores for each cNMF module using Seurat function AddModuleScore. Prior to this, we removed gene redundancy in cNMF modules by assigning each gene to the cNMF module with the highest spectra score ensuring independence when computing module scores and pairwise correlation. To compute pairwise Spearman correlation between cNMF modules across samples, we computed the mean score for each cNMF module across cells of the relative compartment. For the fibroblasts and endothelial cell compartments we computed cNMF modules from metacells, as these showed better performance compared to cNMF modules when using individual cells. We also merged cNMF modules whenever their expression was highly correlated across cells (taking the top 20 genes for each to compute a combined score) and removed others deemed to represent doublets, resulting in a total number of 54 cNMF that can be found in Table S5.

### Copy-number variations inference

We used the package InferCNV (v1.14.2) to infer copy-number variations (CNVs) in the epithelial compartment of the scRNA-seq data. We used a set of normal distal lung cell types including normal mesothelial cells as reference (unpublished). We computed a CNV load score per cell by summing the absolute CNV scores per cell and then normalized the resulting values to the 3^rd^ quantile across cells per sample. A combination of the CNV load distribution and UMAP cell clustering of epithelial cells was used to identify true malignant cells. To infer genomic interaction with cancer cNMF modules, we applied the following strategy: 1) metacells were computed using hdWGCNA function MetacellsByGroups (k=50, max_shared=30) excluding low cell number samples P1 P3 and P13. 2) We selected the most frequent CNV chromosomal rearrangements in our data (at regions: chr1p, chr3p, chr4, chr13, chr14, chr22), which were then used to compute a module score inclusive of all genes contained in each chromosomal region. 3) Malignant cNMF modules were recomputed on metacells excluding all genes that overlapped selected genomic regions. 4) Spearman correlation was computed across metacells for each sample and the median Spearman correlation was used for display in heatmap in Figure 1G. Similarly, for validating this analysis in the Bueno cohort, we took the average expression of the genes in each malignant cNMF module, excluding genes within a CNV region, and computed Spearman correlation with the average expression of genes in each CNV region across samples.

### Bulk RNA-seq datasets acquisition and analysis

RSEM-normalized count matrix including 82 bulk RNA-seq samples as part of the TCGA MESO cohort (Hmeljak cohort) was downloaded using the R package cgdsr (v1.3) of cBioPortal (http://www.cbioportal.org), an online database built for cancer genomics along with metadata including histology information. The Bueno cohort was downloaded from the European Genome-phenome Archive (EGA) under accession number EGAS00001001563 as RPKM-normalized count matrix including 216 bulk RNA-seq samples along with metadata information including histological and molecular subtypes. Both datasets were log2 normalized before any downstream analysis. To compute the score for each scRNA-seq malignant cNMF program we averaged the expression of the top 20 genes for each malignant cNMF module. For cNMF programs identified in other compartments, we first corrected bulk normalized expression for a given cell type abundance (e.g., T cells) by using the function removeBatchEffect from limma R package v3.54.0, where we designated the expression of a canonical marker for a given cell type (e.g., *CD3D* for T cells) as covariate. For immune content correction we used the *PTPRC* marker gene, for T cell infiltration we used *CD3D* and for endothelial content we used *VWF*. This was done to ensure that differences observed in the bulk were not caused by higher abundance of the compartment program assessed. Additionally, to validate T-cell cNMF module enrichment across molecular subtypes, we selected most specific markers for each module: *CD8A, CD8B* for Tm2, *HAVCR2* for Tm5 and *FOXP3 TNFRSF18, ILRA* for Tm7. Similarly, we selected most specific marker genes for the fetal *PLVAP*+ EC subpopulation (*ESM1, PLVAP, TP53I11,* and *INSR*) to compute survival analysis.

### Bulk RNA-seq cell type deconvolution

Cell type deconvolution of bulk RNA-seq samples were performed using the package TED (BayesPrism) (v2.0) (23). BayesPrism is a Bayesian framework that references on cell type expression profiles in scRNA-seq data to statistically estimate the proportion of corresponding cell types in bulk samples. To identify significant differences in the deconvolved cell type proportions we calculated *P* values using the Dirichlet-multinomial regression analysis, implemented by the R package DirichletReg. Since cell compositions sum to one, there is an inversely proportional relationship between cell fractions. Dirichlet-multinomial regression models these dependencies by accounting for the proportions of all other cell subsets when comparing the difference in one cell subset between two PM sample groups (e.g., difference in T cells between sarcomatoid and epithelioid molecular subtypes). Dirichlet regression was used to assess significant variation in cell type abundances from deconvolved bulk-RNA cohorts and scRNA-seq data.

### Assigning bulk RNA-seq based molecular subtypes to malignant single cells

In the Bueno cohort, molecular subtypes were defined based on transcriptomic consensus clustering and assigned to four categories: sarcomatoid, biphasic-S, biphasic-E, and epithelioid (11). Two-dimensional representation of PM subtypes in malignant cells was carried out similarly to Neftel et al (62). Cells were first separated into sarcomatoid/biphasic-S versus epithelioid/biphasic-E by the sign of D = log2(max(sarcomatoid,biphasic-S) – max(epithelioid,biphasic-E)-1), and D defined the y axis of all cells. For sarcomatoid/biphasic-S cells (i.e., D>0), the x axis value was defined as biphasic-S –sarcomatoid and for epithelioid/biphasic-E cells (i.e., D<0), the x axis was defined as epithelioid - biphasic-E.

### Survival analysis

We used a Kaplan-Meier (KM) model to estimate the survival function using the Bueno and Hmeljak cohorts, stratified by their expression levels of various gene modules that can serve as potential prognostic biomarkers. To adjust for histology or molecular subtype groups, we used a stratified Cox proportional hazards regression model and computed *P* values. Both models were implemented using the survival (v.3.4-0) R package. For Kaplan-Meier models we grouped the samples into three groups based on their module score, with high assigned to the first quartile, medium (med) to the second and third quartile and low to the fourth quartile, whereas we used both continuous and stratified values in Cox proportional hazards regression models (reported as (C) *P* value and (S) *P* value respectively). *P* values based on the log-rank test and the chi-square test were used to determine the statistical significance of survival outcomes among the three groups in the adjusted Cox and KM models, respectively.

### SCENIC analysis

SCENIC (v.1.1.2) was run using default settings as described (32) on the myeloid, TAM and endothelial cells. With its implementation in R, SCENIC was run using the 500bp and 10kb motif databases for GENIE3 and RCisTarget. The regulon activity scores (AUC) were calculated using the AUCell (v.4.2) R package for normal and fetal endothelial cells using regulon information from the PM endothelial cells.

### NicheNet analysis

In order to explore the potential regulation mechanisms of modules, we applied the NicheNet package (v2.0.4) implemented in R to predict potential upstream ligands in the TMEs of specific gene signatures. The receiver was defined as the cell population most highly expressing a given module and the sender was the other cell types. Background expressed genes were defined as the intersection of the top 5000 variable features in the receiver cells and the ligand candidates in the ligand-target matrix database provided by NicheNet.

### TCR-seq analysis

TCR analysis was performed using R package scRepertoire (v2.0) (63). Filtered contig lists from each sample outputted from Cellranger were combined using function CombineTCR and mapped to expression data via barcodes using combineExpression function. Clonotypes were labelled as non-expanded, expanded small (n > 1 and n ≤ 5, Small) and expanded large (n > 5, Large). To assess exhaustion in expanded clonotypes we used the computed score for the exhaustion module Tm5 and averaged the score per clonotype across cells. For clonal overlap across samples and across sites we used the ‘CTstrict’ clonecall.

### Immunohistochemistry (IHC)

Paraffin-embedded human mesothelioma tumor samples from all three histological subtypes—epithelioid, biphasic, and sarcomatoid—as well as uninvolved normal lung tissues from lung adenocarcinoma patients, were sourced from the Biorepository tissue bank at the Icahn School of Medicine at Mount Sinai (ISMMS). These tissue samples were procured in accordance with protocols approved by the Institutional Review Board (IRB) of ISMMS. For IHC, 3μm sections of these paraffin-embedded tissue sections were utilized. The IHC process was conducted using the VENTANA Discovery Ultra system (Roche) following the manufacturer’s protocols. This involved de-paraffinization of the tissue sections, followed by sequential staining with primary antibody for CD31 (Roche) and PLVAP (Proteintech). Each primary antibody application was succeeded by the application of corresponding secondary antibodies—DISCOVERY OmniMap anti-Mouse HRP (RUO) Catalog # 760-4310, and DISCOVERY Anti-Mouse HQ Catalog # 760-4814. The signals were then developed using different colors: the DISCOVERY ChromoMap DAB kit (RUO) Catalog # 760-159 for brown and the DISCOVERY Purple kit (RUO) Catalog # 760-229 for purple. Crucially, after each staining phase, slides underwent a process of inhibition, heat denaturation, and neutralization. Subsequently, tissues were counterstained with Hematoxylin to highlight the nuclei in blue. The stained sections were imaged using NanoZoomer S60 Digital slide scanner (Hamamatsu), and the acquired images were analysed using the HALO® Image Analysis Platform (Indica Labs). CD31+ vessels, characterized by brown-stained particles in the cytoplasm, were quantified. Simultaneously, PLVAP-positive cells, discerned by purple-stained particles, were identified. The percentage of CD31 and PLVAP double-positive endothelial cells within blood vessels was then calculated for graphical representation. For each sample, quantification was conducted on 9 to 23 randomly selected regions of interest (ROIs). Statistical significance of the findings was assessed using a paired Student’s t-test.

### Immunophenotyping of mesothelioma cell lines and NK cells

We performed immunophenotyping on all four mesothelioma cell lines used in this study: NCI-H28, MSTO-211H, NCI-H2052, and NCI-H2452. Each cell line was treated overnight in fully supplemented RPMI medium, either with or without 200 ng/ml recombinant human Interferon Gamma (rhIFNγ). Following treatment, cells were stained with Zombie NIR™ (Biolegend) for viability assessment. Subsequently, Fc blocking was performed using TruStain FcX, and the cells were stained with HLAE PE antibody (Biolegend) to assess the surface expression of these receptor. The comprehensive analysis of receptor expression was conducted using flow cytometry with a Cytek Aurora system. Peripheral Blood Mononuclear Cells (PBMCs) were isolated from healthy donor’s blood using density gradient centrifugation with Lymphoprep™ (STEMCELL Technologies) as the separation medium. The freshly isolated PBMCs were subsequently cultured in human NK MACS® medium (Miltenyi Biotec) for 2 to 3 weeks to ensure optimal in vitro expansion of NK cells. NK cells were analyzed by flow cytometry for expression of NKG2A using above mentioned protocol but with anti-human NKG2A PECy5 antibody (Biolegend).

### Mesothelioma-NK Cell Co-culture assay

Mesothelioma cell lines were prepared by incubation with or without 200 ng/ml rhIFNγ in fully supplemented RPMI medium overnight. For the co-culture assay, mesothelioma cell lines were further pre-treated with anti-MHC class I antibody (10 μg/ml) (Clone W6/32, Biolegend) for 1 hour, whereas blood derived invitro expanded NK cells were pre-treated with anti-NKG2A (10 μg/ml) (Beckman Coulter) antibody for 1 hour. Subsequently, NK and mesothelioma cells were combined in a 96-well plate at an effector to target (E:T) ratio of 6:1. Anti-CD107a-BV785 antibody (Biolegend) at 1:500 dilution was added to the cells. After 1 hour, the culture was supplemented with 0.5X concentrations of both brefeldin A and monensin (Biolegend) to facilitate cytokine retention within the cells. The co-cultured cells were then incubated for a total of 16 hours. Post-incubation, the cells underwent staining with Zombie NIR™ (Biolegend) for viability assessment, Fc blocked using TruStain FcX, and then with surface antibodies, including CD45 BUV395, CD3 BUV496, CD4 BV570, CD8 PerCP Cy5.5, CD56 BUV805, PD-1 BV711, and NKG2A PECy5 (Biolegend). The cells were fixed using IC fixation buffer (Biolegend) and intracellularly stained using granzyme A AF700 and IFNγ PE antibodies in 1x Permeabilization Buffer (Biolegend). Finally, the stained samples were subjected to flow cytometric analysis using a Cytek Aurora system to quantitatively assess NK cell degranulation and cytokine production. Flow Jo was used for flow cytometry data analysis. We employed the Flow AI algorithm via the FlowJo software platform. This approach facilitated the automated identification and exclusion of aberrant events, ensuring high-quality data for subsequent analysis. The parameters and thresholds for Flow AI were set in accordance with the software’s guidelines to optimize data integrity and analytical accuracy.

## Data availability

All scRNA-seq, CITE-seq and TCR-seq data have been deposited in the GEO and are available under accession number GSE190597.

## Acknowledgements

We thank all members of the Mount Sinai Genomics Core, Human Immune Monitoring Core, and ImmunAI that helped with single cell profiling and sequencing of tumor and blood samples, respectively. This research was also conducted with support from the Biorepository and Pathology Core at Mount Sinai. This work was supported in part by American Association for Thoracic Surgery/Women in Thoracic Surgery Mid-career Investigator Grant for A.W. and ImmunAI and ISMMS seed funding for A.M.T.

## Author contributions

A.M.T. conceived the project and designed the study. B.G., P.C., W.Z., G.K., and A.B. performed computational analyses and A.M.T., B.G., P.C. interpreted the results. A.W., T.M., and R.F. performed surgical resections and provided scientific feedback along with B.F. and S.H. K.D., R.Sw., and W.Z. acquired and processed PM tissue. R.Se., K.B., S.S., E.K., T.D., R.C., S.K.S. and Z.C. performed and supervised 10x loading, library construction, and sequencing. M.M. set up lung tissue acquisition pipeline and supervised R.Sw. along with S.G. A.M.T., A.T., E.M., A.H., and K.D. designed the co-culture experiment, performed by K.D and E.M. A.M.T., M.G., J.A., A.S., R.B. identified and processed PM tissue sections for IHC that were analyzed by J.A., M.G. and K.D. B.G., K.D., P.C., and A.M.T. wrote the manuscript with input and approval from all authors.

---

**Table S1** Additional clinical information for each patient in the PM single cell cohort.

**Table S2** *De novo* markers discovered for each major cell type annotation in tumor samples.

**Table S3** *De novo* markers discovered for each major cell type annotation in the peripheral blood samples.

**Table S4** *De novo* markers discovered for each detailed cell type annotation in the peripheral blood samples.

**Table S5** Gene signatures for all 54 *de novo* discovered PM cNMF modules.

**Table S6** SCENIC-predicted regulon targets of ETS1 and MEF2C in PM endothelial cells.

**Figure S1.**
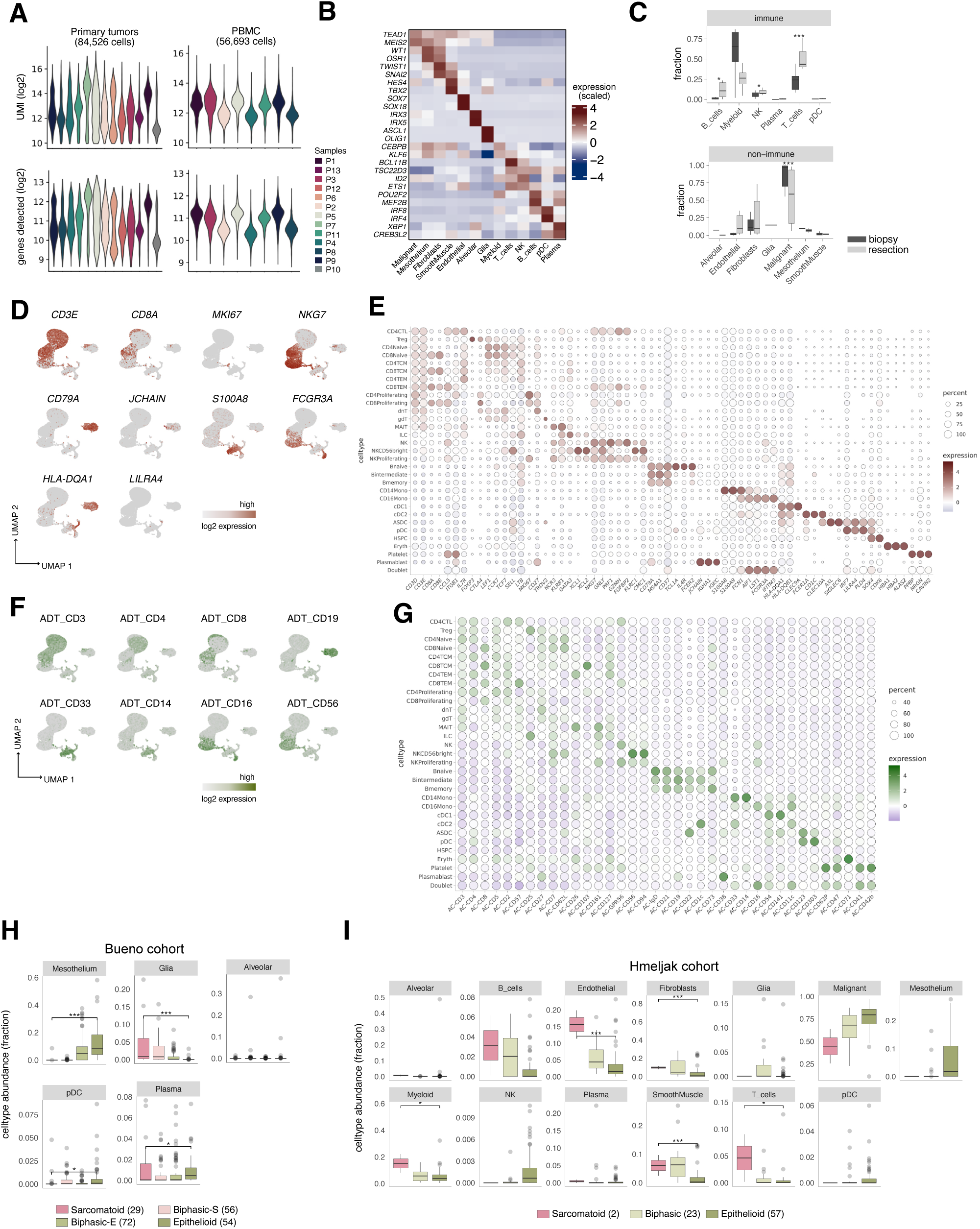
Single-cell catalogue of PM tumor and PBMC samples. **A,** Violin plot of key quality control metrics of the tumor (left) and PBMC (right) scRNA-seq data per patient. **B**, Top transcription factor (TF) makers per cell type annotation in the primary tumors. **C**, Distribution of cell type abundances split by immune (CD45+) and non-immune (CD45-) compartment and sampling procedure (resection versus biopsy). FDR-adjusted *P* values were determined by Dirichlet-multinomial regression model. **D, F**, Feature plots of key RNA **(D)** and protein **(F)** markers in the PBMC data. **E, G**, Dot plots of top *de novo* discovered RNA **(E)** and protein markers **(G)** per cell type annotation. **H, I,** Sample distributions based on fraction of cell types and grouped by molecular subtype in the Bueno cohort (**H**) and grouped by histology in the Hmeljak cohort (**I**). FDR-adjusted *P* values comparing difference between sarcomatoid and epithelioid subtypes were determined by Dirichlet-multinomial regression model that takes into account dependencies in proportions between cell types. TCM = central memory T cells; TEM = effector memory T cells; dnT = double negative T cells; MAIT = mucosal associated invariant T cells; Eryth = erythrocytes; CTL = circulating T lymphocytes; gdT = gamma delta T cells; HSPC = hematopoietic stem and progenitor cells; ILC = innate lymphoid cells; ASDC = AXL^+^ SIGLEC6^+^ dendritic cells. *p<0.05, **p<0.01, ***p<0.001, ****p<0.0001.

**Figure S2.**
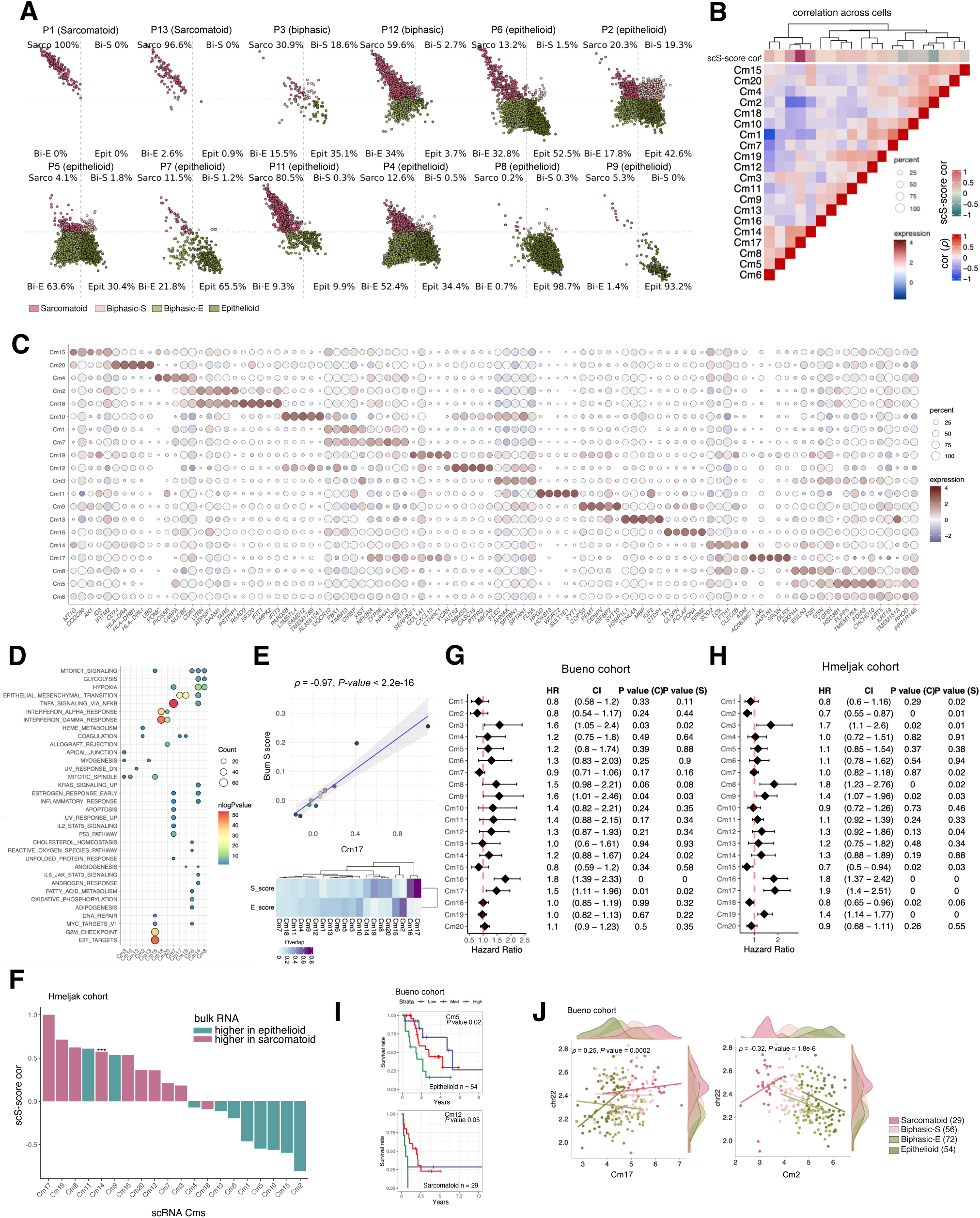
Unbiased discovery of PM cancer programs and association with patient survival. **A,** 2D visualization of malignant cells scored using the four molecular subtypes from Bueno et al for each sample. Clinical histology of the cancers upon diagnosis are reported in parentheses (Methods). **B**-**C,** Pairwise Spearman correlations (**B)** across all cells for the 20 Cms identified in malignant cells. Dot plot (**C)** showing the expression and percentage of cells expressing top markers for each Cm. **D**, Pathway enrichment analysis showing the 10 most enriched pathways for each Cm within the HALLMARK gene categories. Only Cms with significant categories are shown. Dot size represents number of genes in the category and color represent −log10 (*P* value). **E,** Top: Spearman correlation and *P* value across scRNA-seq samples between mean Cm17 module score (scS-score) and mean Blum et al S_score (top 20 genes) in malignant cells. Bottom: Fraction overlap in genes between the markers for each of the 20 malignant Cms discovered in our scRNA-seq cohort and the two gene sets associated with a sarcomatoid subtype (S-score) and epithelioid subtype (E-score) from Blum et al. **F,** Cms ranked by correlation to the scS-score and colored by their enrichment for either epithelioid (green) or sarcomatoid (red) histological subtypes from the Hmeljak cohort. FDR-adjusted *P* values were computed using Welch Two Sample *t*-test. *p<0.05, **p<0.01, ***p<0.001, ****p<0.0001. **G, H,** Cox proportional hazard regression models for all the 20 Cms using survival information from the Bueno cohort (**G**) and the Hmeljak cohort (**H**). **I,** Kaplan-Mayer survival curves of Cms significantly impacting survival within molecular subtypes in the Bueno cohort. **J,** Spearman correlation and *P* values of samples in the Bueno cohort using average expression of all genes residing on chr22 versus average expression of genes included in Cm17 (left) and Cm2 (right).

**Figure S3.**
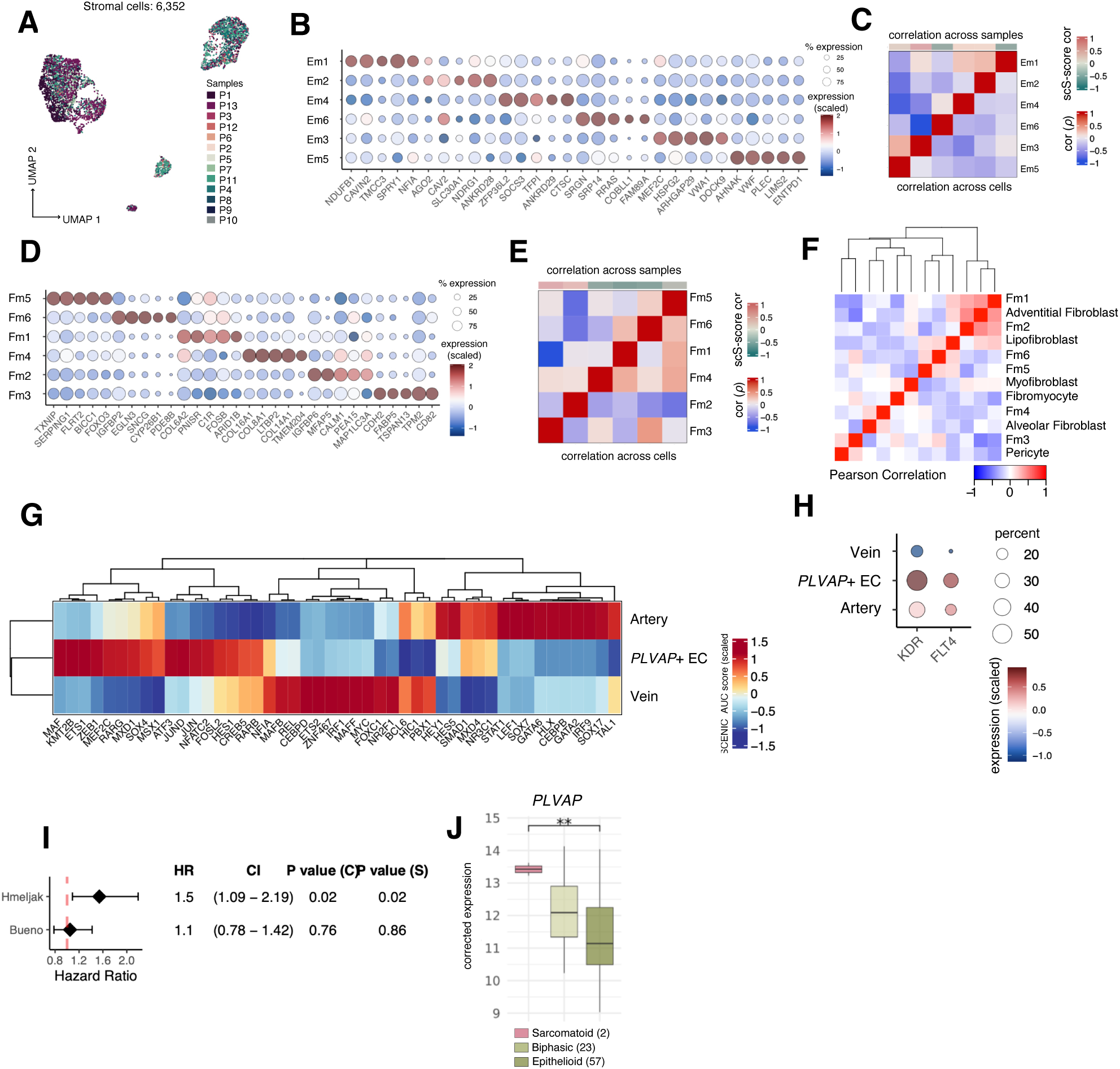
Fetal-like, cancer-specific PLVAP+ endothelial cells associate with angiogenesis. **A,** UMAP of integrated tumor endothelial and mesenchymal cells colored by sample. **B-C**, Dot plot showing the expression and percentage of cells expressing top markers in the endothelial gene modules (Ems) (**B**) and their pairwise Spearman correlation across samples (above diagonal) and cells (below diagonal) **(C). D**, Dot plot showing expression and percentage cells expressing top markers in the CAF gene modules (Fms). **E**, Heatmap of the pairwise Spearman correlation between Fms across samples (above diagonal) and cells (below diagonal). **F**, Heatmap of pairwise Pearson correlation coefficient between the Fms and normal lung mesenchymal cell subset average expression profiles (29). **G**, Heatmap of the average TF regulon activities inferred by SCENIC for each endothelial cell subset. **H**, Dot plot showing expression and percentage cells expressing VEGFA receptor genes *KDR* and *FLT4* for each EC subset. **I**, Cox proportional hazard regression analysis (adjusted for molecular subtype in the Bueno cohort and histology in the Hmeljak cohort) based on the expression of fetal-like PLVAP+ EC subset marker genes (Methods) and survival information from the and Hmeljak (top) and Bueno cohorts (bottom). **J**, Sample distributions based on *PLVAP* expression in Hmeljak cohort after correction for endothelial content. *P* values were computed using Welch Two Sample *t*-tests. **p<0.01.

**Figure S4.**
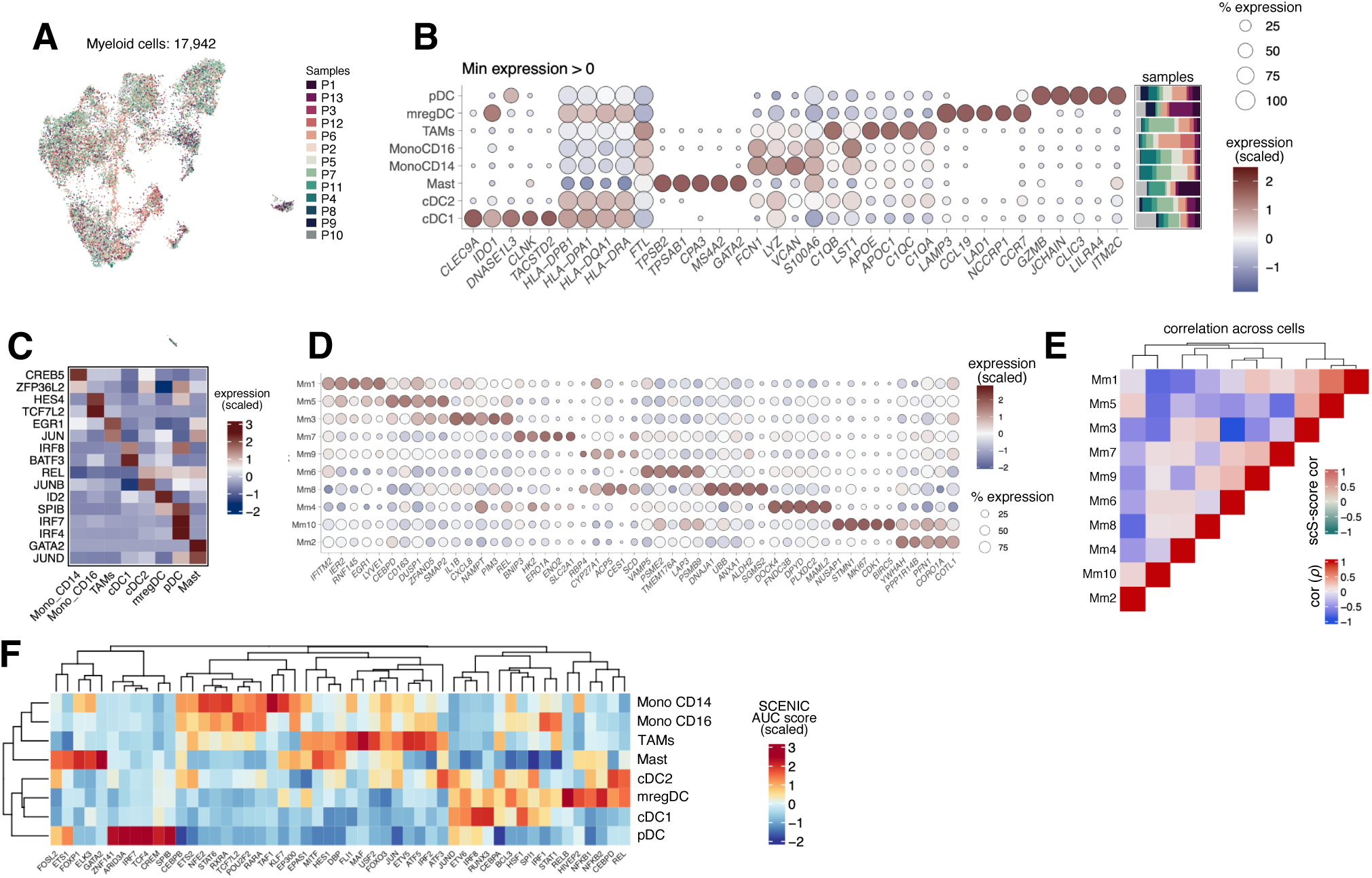
Macrophages in scS-high PM express *CXCL9/10/11* and likely contribute to T-cell infiltration. **A**, UMAP of integrated tumor myeloid cells colored by sample. **B**, Dot plot showing the expression and percentage of cells expressing top markers for each myeloid subset and relative sample composition (right, stacked bar plots). **C**, Top transcription factor (TF) markers per myeloid cell subset. **D,** Dot plot showing the expression and percentage of cells expressing top markers in the TAM gene modules (Mms). **E**, Heatmap of the pairwise Spearman correlation between Mms across cells. **F,** Heatmap of the SCENIC significant regulon activities (scaled AUC score) and corresponding TFs (columns) in each myeloid subtype.

**Figure S5.**
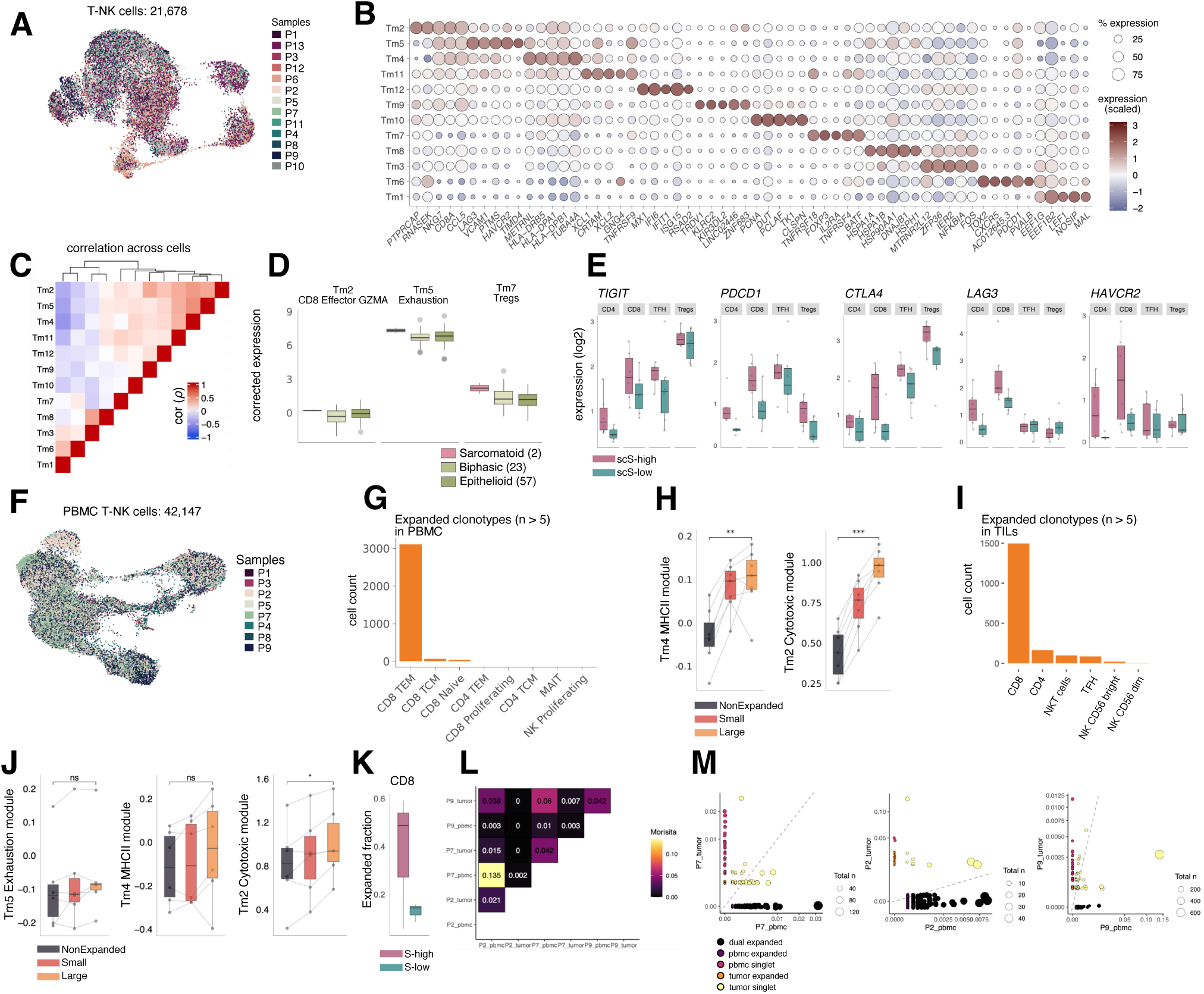
Molecular dissection of T cell programs and IC molecules shows association with scS-score. **A,** UMAP of integrated T and NK cells in the primary tumor colored by sample. **B,** Dot plot showing the expression and percentage of cells expressing top markers in the T cells modules (Tms). **C**, Heatmap of the pairwise Spearman correlation between Tms across cells. **D**, Sample distributions of the Tm2, Tm5, and Tm7 marker expression in the Hmeljak cohort for each PM histological subtype after correcting for immune content. FDR-adjusted *P* values were computed comparing sarcomatoid and epithelioid subtypes using Welch Two Sample *t*-test. **E**, Sample distribution of the log2-normalized expression of key IC molecules for each T-cell subset split by scS-high and scS-low samples. **F,** UMAP of integrated T and NK cells in PBMC and colored by sample. **G**, Number of cells with expanded clonotypes (large) across PBMC T-NK subsets. **H,** PBMC sample distribution of the Tm4 and Tm2 mean module scores in non-expanded vs expanded (small and large) CD8 T cells. I, Number of cells with expanded clonotypes (large) across tumor cell subsets. **J,** Tumor sample distribution of the Tm5, Tm4, and Tm2 mean module scores in in non-expanded vs expanded (small and large) CD8 T cells. *P* values were computed using Welch Two Sample *t*-test. **K,** Fraction of expanded clonotypes in CD8 T cells in tumor samples with detectable TCR sequences split by scS-high and scS-low samples. **L**, Overlap of TCR clonotypes as computed by Morisita score in 3 patients with available scTCR-seq data from both tumor and blood. **M,** Expanded clonotypes identified both in tumor and PBMC for the three patients with matching TCR-seq data available. *p<0.05, **p<0.01, ***p<0.001, ****p<0.0001.

**Figure S6.**
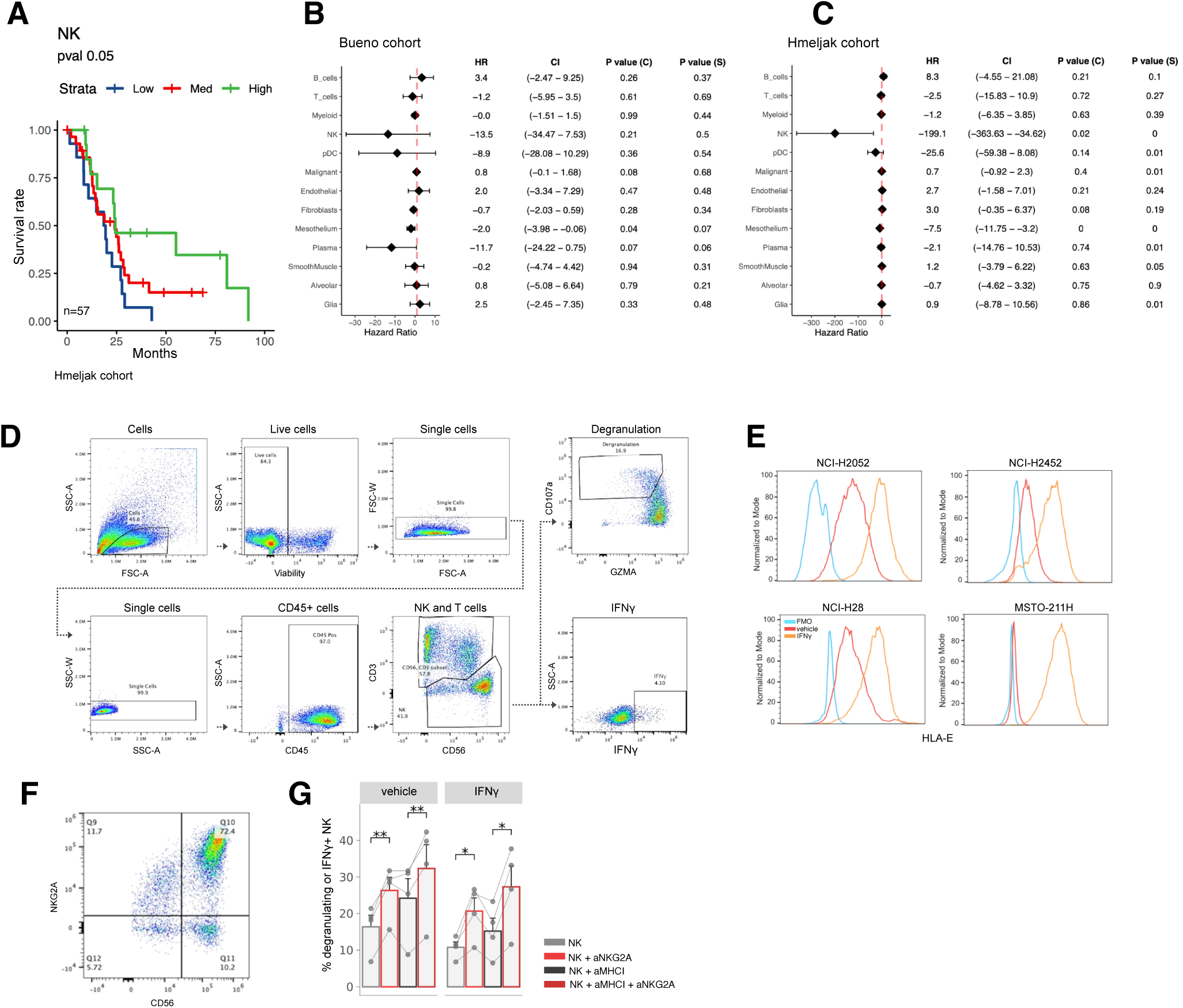
NK cell IC blockade targeting NKG2A as a novel therapeutic strategy in PM. **A,** Survival curve stratifying NK cell abundance (deconvolved using BayesPrism) in epithelioid samples in the Hmeljak cohort. **B-C,** Cox proportional hazard regression models for all the predicted cell types (deconvolved using BayesPrism) using survival information from the Bueno cohort (**B**) and the Hmeljak cohort (**C**). **D,** Flow cytometry gating strategy for the cancer-NK cell co-culture experiments. **E,** Expression of HLA-E on 4 mesothelioma cell lines with and without interferon gamma stimulation. **F,** NKG2A expression on *in-vitro* expanded NK cells. **G,** Bar graph displaying the percentages of NK cells either producing IFNγ or undergoing degranulation (Boolean gating). *P* values were computed using a paired Student’s t-test and error bars represent standard error. *p<0.05, **p<0.01, ***p<0.001, ****p<0.0001.

